# Circadian oscillation of perireceptor events influence olfactory sensitivity in diurnal and nocturnal mosquitoes

**DOI:** 10.1101/2023.10.19.563057

**Authors:** Tanwee Das De, Julien Pelletier, Satyajeet Gupta, Madhavinadha Prasad Kona, Om P. Singh, Rajnikant Dixit, Rickard Ignell, Krishanpal Karmodiya

## Abstract

Olfaction and circadian rhythm gate different behaviors in mosquitoes that are important for their capacity to transmit disease. However, the mechanisms of odor detection, and the circadian-guided changes in olfactory sensitivity across different mosquito species, remain largely unexplored. To this end, we performed a circadian-dependent RNA-sequencing study of the peripheral olfactory- and brain tissues of female *Anopheles culicifacies* and *Aedes aegypti* mosquitoes. Data analysis revealed a significant upregulation of genes encoding: (a) odorant binding proteins (OBPs), required for transportation of odorant molecules towards the olfactory receptors, and (b) xenobiotic-metabolizing enzymes (XMEs) during the day time in *Aedes aegypti* and during the dusk-transition phase in *Anopheles culicifacies*. While XMEs primarily function in the elimination of toxic xenobiotics, concurrent elevation of XMEs and OBPs are hypothesized to act cumulatively to regulate perireceptor events and odorant sensitivity. Electroantennographic analysis with both *Anopheles gambiae* and *Aedes aegypti* against diverse behaviorally relevant odorants, combined with XMEs inhibitors and RNA interference, establish the proof-of-concept that XMEs function in perireceptor events during odorant detection and influence the odorant sensitivity in mosquitoes. Additionally, the RNA-sequencing and RNAi-mediated knockdown data revealed that daily temporal modulation of neuronal serine proteases may facilitate the consolidation of the brain function, and influence the odor detection process in both diurnal and nocturnal mosquitoes. These findings provide the impetus to further explore the species-specific rhythmic expression pattern of the neuro-olfactory encoded molecular factors, which could pave the way to develop and implement successful mosquito control methods.

**Graphical Abstract:** 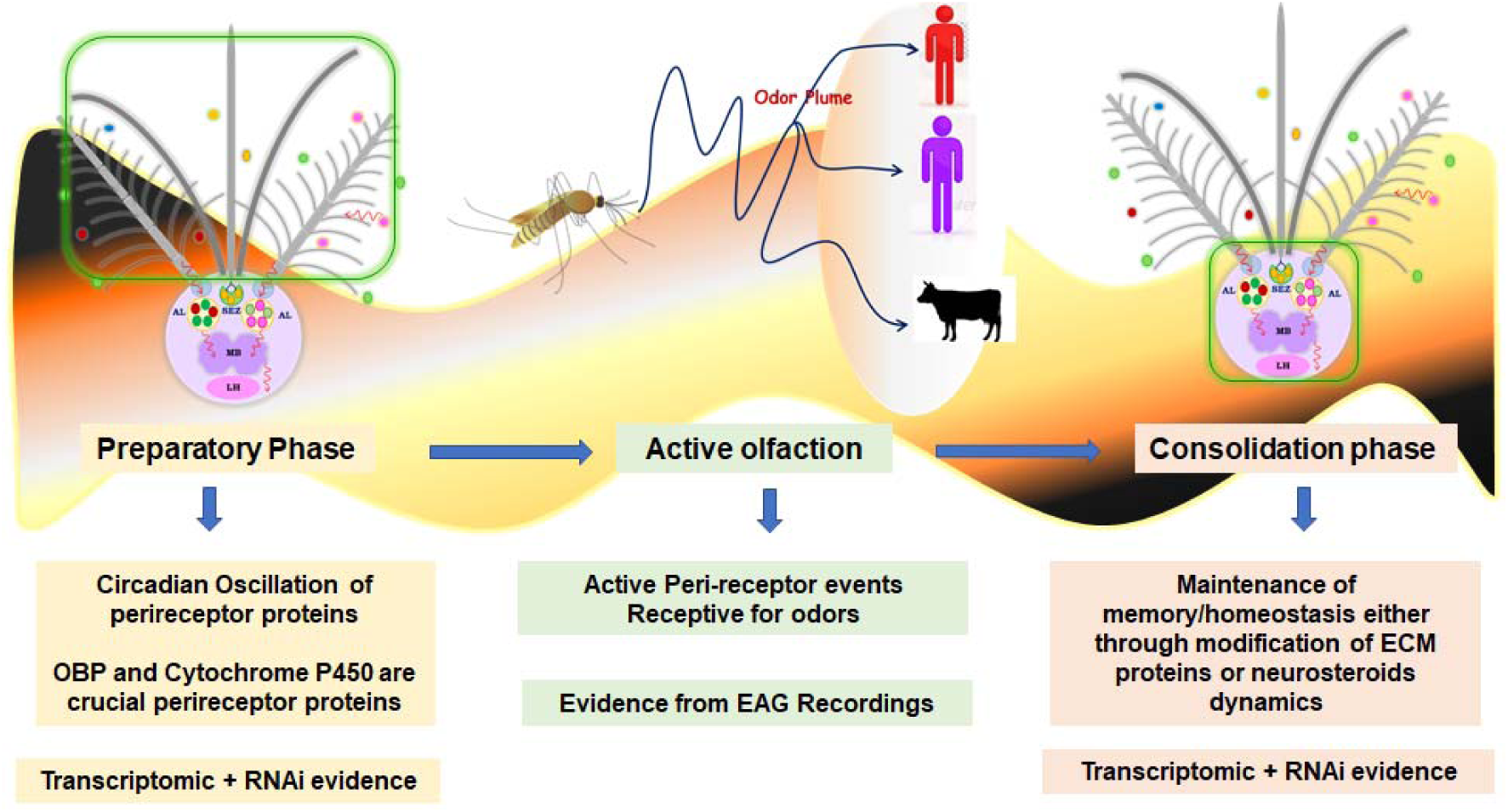

**Highlights:**

- Circadian oscillation of perireceptor proteins possibly influences time-of-day dependent change olfactory sensitivity in diurnal and nocturnal mosquitoes
- Diurnal and nocturnal mosquitoes depict distinct dynamic change in perireceptor proteins
- Inhibition of cytochrome P450 gene minimizes antennal response to different odorants
- Neuronal serine protease may consolidate brain function and odor detection

## Introduction

Mosquitoes are vectors of many infectious diseases and pose a serious-threat to society. The lack of effective vaccines against mosquito-borne diseases underscores the importance of mosquito control approaches (*1*). However, insecticide resistance, its adverse environmental impact, and the concurrent adaptive evolution of vector populations highlight the need to formulate ideas for developing novel vector control strategies (*2*, *3*). Notably, olfaction is prerequisite for feeding, mating, and survival in mosquitoes, in which olfactory learning and memory may modulate these behavioral activities, affecting the fitness of these insects (*4–6*). Furthermore, different species of mosquitoes exhibit plasticity and distinct rhythms in their daily activity pattern, including locomotion, feeding, mating, blood-feeding, and oviposition, facilitating their adaptation into separate time-niches (*7*, *8*), but the underlying molecular mechanism for the heterogenous temporal activity remains to be explored.

During the evolution from cyanobacteria to higher vertebrates, internal biological clocks have facilitated the ecological adaptation of organisms, by temporal regulation of the sleep-wake cycle, behavior, and cognitive performance (*9–11*). The circadian clocks are governed by the endogenous molecular oscillators, which autonomously synchronize external environmental cues, such as light and temperature, broadly known as zeitgeber, with the internal physiological events (*9*). The core circadian oscillators are comprised of several clock genes, *e.g., clock (clk), cycle (cyc), period (per), timeless (tim), pigment dispersing factor (pdf),* and *cryptochrome 2 (cry2)*, that regulate the circadian rhythm via two interlocked transcription/translation negative feedback-loops (*11–13*). Several studies in *Drosophila* have demonstrated that the pacemaker of the circadian clock is present within a set of neurons in the brain, which entrain and transmit zeitgeber information towards the downstream peripheral clocks (*11*, *14*). A comparative study between the neural circuits of diurnal and nocturnal mosquitoes depicts similar but distinct organization of neuronal circuits, and the molecular clock is found to be regulated by anti-phasic expression of clock-proteins ensuring their diurnality and nocturnality (*8*, *15*). But, how the pacemaker differentially influences peripheral clock activity present in the olfactory system and modulates olfactory sensitivity has not been studied in detail.

Odor coding in insects involves the combinatorial activation of olfactory receptors, including odorant receptors (ORs) and ionotropic receptors (IRs), and transmission of olfactory information via olfactory sensory neurons (OSNs) towards the higher olfactory centers in the brain that ultimately results in an appropriate behavioral response (*16*, *17*). For the activation of olfactory receptors, the odorant molecules need to traverse through the aqueous interphase of the sensillar lymph (*18*). Any biochemical reactions that alter the chemical nature and binding property of the odorants in the lymph may have an impact on olfactory sensitivity and selectivity (*19*, *20*). These initial physiological events are called the “perireceptor events”, and bridge the gap between the chemical environment and the olfactory receptors, thereby regulating the peripheral detection of the olfactory signals (*19*, *21*). Primarily two groups of proteins are present in the lymph surrounding the sensory neurons: (a) odorant-binding proteins of two classes, the odorant-binding proteins (OBPs) and the chemosensory proteins (CSPs), and (b) xenobiotic metabolizing enzymes (XMEs), which affect odorant entry and clearance, respectively (*19*). The OBPs and CSPs reversibly bind with the odorants and deliver them to the olfactory receptors (*19*, *22*). The XMEs, including cytochrome P450 (CYP450: phase I XMEs), UDP-glucosyl transferase (UGT: phase II XMEs), and efflux transporters (phase III XMEs), may either introduce a functional group to the hydrophobic odorant molecules to make them more hydrophilic for facilitating their transportation, and thus modulate olfactory sensitivity, or inactivate the odorant by biotransformation or degradation, and thus avoid over-sensitization (*19*, *23*, *24*). Therefore, the cumulative actions of OBPs/CSPs and XMEs may directly affect the residence time of the odorants for receptor activation, and thereby influence olfactory detection.

Although spatio-temporal expression patterns of olfactory genes are under the control of hard-wired genetic systems, evidence from previous studies indicates that changes in environmental conditions, and the internal nutritional status have a strong impact on their expression (*15*, *25*, *26*). For example, blood-meal ingestion, circadian cycle, and aging cause transcriptional switching of the olfactory encoded molecular factors (*e.g.,* ORs, IRs, OBPs, CSPs) that lead to a change in the olfactory sensitivity of mosquitoes (*25–30*). A subset of OBPs and CSPs show more dynamic and transient expression patterns according to changing internal physiological conditions (*25*, *26*, *30*). In contrast, ORs show highly selective and programmed expressions that are modulated predominantly by age and experience-guided events (*28*, *29*). Previous studies by Rund *et al.* also indicated that OBPs display more pronounced rhythmic expression compared to ORs, and that the peak abundance of a subset of OBPs at night coincides with higher olfactory sensitivity in *Anopheles gambiae* to host-derived odorants (*25*). Although several detoxification categories of genes (XMEs) displayed circadian rhythmicity in the heads of both diurnal and nocturnal mosquitoes (*26*, *30*, *31*), their function in odorant discrimination and degradation processes have so far only been tested in the fruit fly and beetle (*23*, *24*).

In vertebrates and invertebrates, it is well documented that circadian phase-dependent training can influence olfactory memory acquisition and consolidation of brain functions (*32*–*34*). However, the acquired memories are very labile and change by alternation of external and internal factors (*32*, *35*). To make the memory stronger and more resilient, cell and system level consolidation are required that reorganize the communication among neurons (*36*, *37*). The synapse and the peri-synaptic structure, comprising pre-and postsynaptic neurons, and the surrounding adhesion molecules, govern the transmission of signals between neurons (*38*, *39*). Circadian-training-induced activation of MAP kinase and AMP-kinase activity, and modulation of downstream gene transcription, may cumulatively influence synaptic plasticity and memory in invertebrates (*40–42*). Previous studies showed that synaptic plasticity and memory are significantly influenced by the strength and number of synaptic connections (*43*, *44*). Furthermore, an increasing number of studies in higher organisms indicate that synaptic extracellular matrix (ECM) has an essential role in signal transmission, reorganizing synaptic framework as well as plasticity (*38*, *39*). A proteolytic cleavage of these molecules is thought to be prerequisite for ECM remodeling, the release of synaptic vesicles, stabilization, and maintenance of synapse, which may cumulatively influence synaptic plasticity (*38*, *39*). Among the several proteases, matrix metalloprotease, serine protease, and aspartic proteases have been recognized as potential neuronal proteases, affecting neuronal plasticity (*39*, *45*). Different mosquito species exhibit environmentally-guided pronounced plasticity in their locomotion, host-seeking, mating, and blood-feeding activities (*46*, *47*). Thus, it can be postulated that the concatenated effects of both olfactory and neuronal plasticity, as well as memory, are the major factors to regulate behaviors of mosquitoes in a given ecological and evolutionary context. But, how the brain of mosquitoes entrains circadian inputs and modulates transcriptional responses that consequently contribute to remodel plastic memory, is unknown.

In order to trace the circadian-guided molecular rhythm in the olfactory and brain tissues, which differentially impact the behavioral responses in diurnal and nocturnal mosquitoes, we conducted a comparative RNA-Seq analysis of *Anopheles culicifacies* and *Aedes aegypti*. Through detailed functional annotation and differential gene expression analysis coupled with RNAi and electroantennographic (EAG) studies on *Anopheles gambiae* and *Ae. aegypti*, we demonstrate that: (i) limited clusters of genes display daily rhythmic expression in both olfactory and brain tissues; (ii) the predominant temporal changes of perireceptor proteins influence odor detection and olfactory sensitivity; and (iii) neuronal serine protease may have important functions in consolidating brain activity and olfactory sensitivity.

## Results

### Temporal differences in transcript abundance in *Anopheles* compared with *Aedes*

To decipher the molecular mechanism of the peripheral olfactory and brain systems influencing species-specific, circadian-dependent olfactory sensitivity, *An. culicifacies* and *Ae. aegypti* were selected as the reference for nocturnal and diurnal species, respectively. Neuro-olfactory tissues were sampled at 6 h intervals, starting from ZT0 to ZT18, from both species for subsequent RNA sequencing (**Figure 1A**). To visualize the effect of the circadian clock on the global transcriptional expression pattern, a normalized read count-based heatmap (**Figure 1B**) and density plot **(Figure S1)** analysis were performed. Both species demonstrated minor time-of-day dependent shifts in transcriptional responses of peripheral olfactory and brain genes (**Fig. 1B** and **Figure S1)**. To corroborate the data with previous observations (*15*, *25*, *26*), the expression pattern of key endogenous regulators of the biological clock was analyzed (**Figure 1C**). As reported earlier, a significant upregulation of *period* and *timeless* during ZT12-ZT18 was observed in both species (**Figure 1C**). Next, the distribution of peak transcriptional changes in both *An. culicifacies* and *Ae. aegypti* was assessed through differential gene-expression analysis. Noticeably, *An. culicifacies* showed a higher abundance of differentially expressed olfactory genes (**Figure 1D**) compared to *Ae. aegypti* (**Figure 1E**) at each time point, when considering ZT0 as the reference time, as both mosquito species remain inactive during the early morning (*50*, *51*). Furthermore, *An. culicifacies* demonstrated maximum enrichment in the number of regulated genes during the dusk transition phase (ZT12), in both peripheral olfactory and brain tissues (**Figure 1D** and **Fig. S2A)**. Similar to previous studies (*26*), the expression of a limited number of rhythmic genes was visualized in *Ae. aegypti,* peaked during ZT12 - ZT18 in the peripheral olfactory system (**Figure 1E**), and during the dusk transition phase (ZT12) in the brain **(Figure S2B)**.

**Figure 1:**
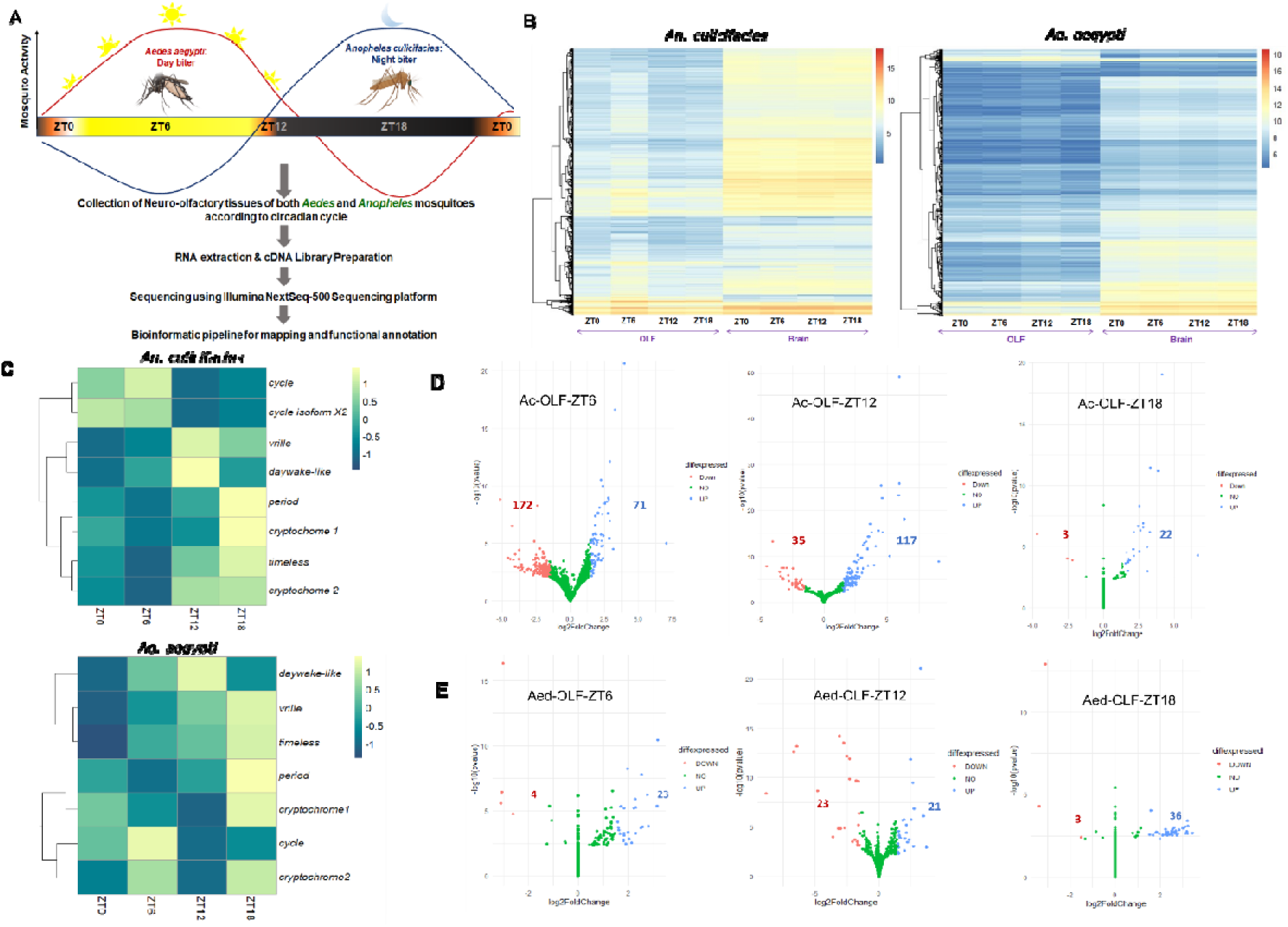
Temporal difference in global transcriptome profile. (A) Technical and experimental workflow for transcriptome analysis of peripheral olfactory and brain tissue. ZT0 (6am) - end of the dawn transition, ZT6 (12pm) - noon, ZT12 (6pm) - time of lights off, ZT18 (12am) - midnight. (B) Normalized read-count-based heatmaps represent the global transcript expression pattern of both peripheral olfactory and brain tissues at all time points. (C) Normalized read-count-based heatmap of central clock genes in the brain of both *An. culicifacies* and *Ae. aegypti*. (D) Volcano plots of differentially expressed genes in the peripheral olfactory system of both *An. culicifacies* and *Ae. aegypti*.

Taken together, the data suggests that the nocturnal *An. culicifacies* may possess a more stringent circadian molecular rhythm in peripheral olfactory and brain tissues.

### Circadian cycle differentially and predominantly expresses olfaction-associated detoxification genes in *Anopheles* and *Aedes*

To analyze the central circadian clock regulated downstream molecular mechanism, which governs diurnality or nocturnality, a detailed gene-set-enrichment analysis was performed. The upregulated genes in the olfactory tissue of *An. culicifacies* were predominantly represented under the functional categories of UDP-glucosyltransferase activity, small molecule metabolic process, oxidoreductase, and monooxygenase activity, as well as ion binding and extracellular region (**Figure 2A**). The downregulated genes were constituted with cytoskeletal motor activity, microtubule-based process during ZT6, and fatty-acid metabolic process during ZT12 **(Figure S3A)**. Notably, the gene-ontology analysis revealed a gradual increase in a subset of oxidoreductase and monooxygenase activity-related genes during ZT6 to ZT12 in *An. culicifacies* (**Figure 2A**). Detailed functional annotation analysis of the oxidoreductase family suggested the predominant occurrence of CYP450 **(Figure 2B, S3B)**, dehydrogenase/reductase SDR family member 11 precursor and prophenoloxidase genes during ZT6-ZT12 **(Supplementary File 1)**. The pathway prediction analysis indicated the notable enrichment of the xenobiotic/drug metabolism pathway and steroid hormone biosynthetic pathways during ZT12, the linoleic acid and arachidonic acid metabolism pathways, as well as the amino-acid biosynthetic pathway during ZT18 in *An. culicifacies* **(Figure. S3C)**. Similar to *An. culicifacies*, the overrepresented upregulated genes of *Ae. aegypti* belong to the functional categories of the regulations of gene expression, biological process as well as oxidoreductase and monooxygenase activity **(Figure 2C, S3D)**. The downregulated genes of *Ae. aegypti* did not show any functional categories probably due to the limited transcriptional change. A maximum enrichment of oxidoreductase and monooxygenase activity-related genes was observed at ZT6 in *Ae. aegypti* (**Figure 2D**), comprising of prophenoloxidases and CYP450 genes (**Supplementary File 1)**. Regulation of gene expression and biological process categories majorly consists of genes encoding transcription factors (GATA and tcf) **(Supplemental File 1)**.

**Figure 2:**
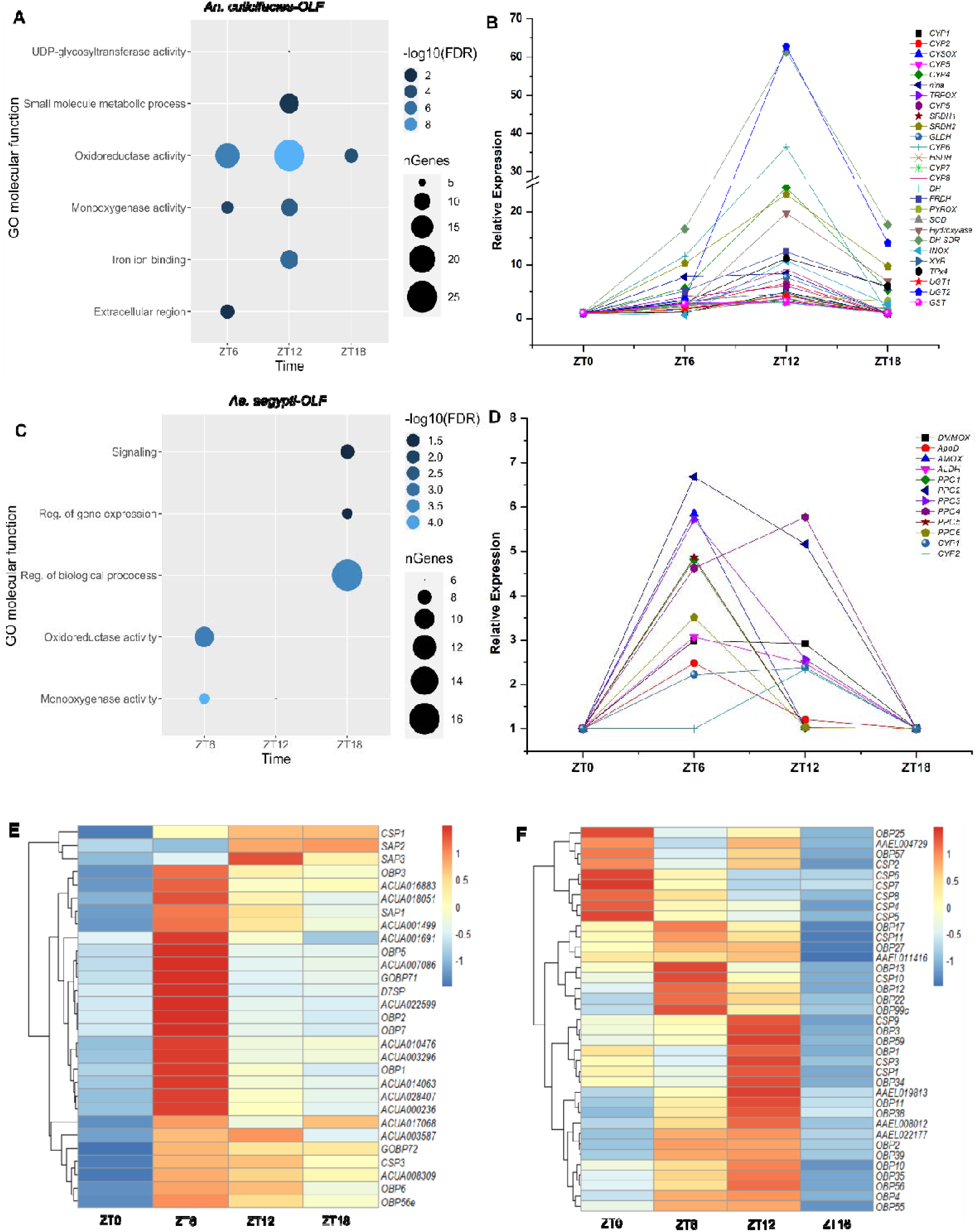
Functional annotation of the circadian-dependent olfactory transcriptome of nocturnal and diurnal mosquitoes. (A) Gene Ontology analysis of upregulated transcripts (>=log2 1.5-fold) in the peripheral olfactory system of *An. culicifacies* when ZT0 is considered as the reference time point. (B) Relative expression profiling of xenobiotic metabolizing pathway-related genes in *An. culicifacies*. CYP: Cytochrome P450; CYSOX: Cysteine dioxygenase; nina: Neither inactivation nor afterpotential protein G; TRPOX: Tryptophan dioxygenase; SRDH: Sorbitol dehydrogenase; GLDH: L-gulonate 3-dehydrogenase; HSDH: hydroxysteroid dehydrogenase; FRDH: farnesol dehydrogenase; PYROX: 4-hydroxyphenylpyruvate dioxygenase; SOD: Superoxide dismutase; DH-SDR: Dehydrogenase/reductase SDR family member 11 precursor; INOX: inositol oxygenase; XYR: D-xylulose reductase A; TPx4: Thioredoxin peroxidase 4; UGT: UDG-Glucosyl transferase. (C) Gene ontology analysis of upregulated transcripts (>=log2 1.2-fold) in the peripheral olfactory system of *Aedes aegypti* when ZT0 is considered as the reference time point. (D) Relative expression profiling of the detoxification category of genes in *Ae. aegypti* according to the circadian clock. DMMOX: Dimethylaniline monooxygenase; ApoD: Apolipoprotein D; AMOX: Amine oxidase; ALDH: Aldehyde dehydrogenase; PPO: prophenoloxidase; CYP450: Cytochrome P450. (E) Normalized read count-based heatmap of odorant binding protein (OBP) genes of *An. culicifacies*. CSP: Chemosensory protein; SAP: Sensory appendages protein. (F) Normalized read count-based heatmap of odorant binding protein (OBP) genes of *Ae. aegypti*. CSP: Chemosensory protein.

Though distinct time-of-day dependent transition peaks of the oxidoreductase category of genes were observed for both *An. culicifacies* and *Ae. aegypti* (ZT12 for *An. culicifacies* and ZT6 for *Ae. aegypti*) (**Figure 2B, D**), a GO-enrichment analysis was unable to track any change in the response-to-stimulus or odorant binding category of genes (including OBPs, CSPs, and olfactory receptors). However, functional annotation of differentially expressed genes revealed a significant upregulation of a subset of CSPs (∼ 5-fold) and OBP6 (∼3.3-fold) transcripts during ZT12 in *An. culicifacies* **(Supplementary File 1).** Olfactory receptor genes did not fulfill the selection criteria (normalized read-count ≥ 50) and were thus excluded for further analysis. For a comprehensive understanding of the rhythmicity of all annotated OBP genes, a normalized read count-based heatmap analysis was conducted for *An. culicifacies* (**Figure 2E**). A tight rhythmic expression of almost all OBPs was observed, with maximum enrichment during ZT6-ZT12 (**Figure 2E**), coinciding with the XMEs expression pattern **(Figure 2B, S3B)**. In contrast, three different clusters of OBP genes in *Ae. aegypti* showed a time-of-day dependent distinct peak in expression starting from ZT0-ZT12 (**Figure 2F**). Notably the diurnal and nocturnal mosquitoes exhibit temporal segregation of minimal olfactory response (*50*, *51*) that are correlated by the suppression of OBP genes transcription at ZT0 in *Anopheles culicifacies* and at ZT18 in *Aedes* aegypti **(Figure 2E, F)**. To assess the role of clock regulation of gene enrichment in *An. culicifacies*, potential clock-light responsive elements were analyzed, including *e.g.,* the E-box element, in the 5kb 5C upstream region from the transcription start-site of the respective upregulated CYP450 and CSP genes. At least five CYP450 genes and five CSPs genes of *An. culicifacies,* possess the multiple consensus sequence of the E-Box element (CACGTG) within their upstream region **(Supplemental File 2).** Similarly, prophenoloxidase, serine protease, and the CYP450 genes of *Ae. aegypti* was observed to consist of at least one conserved E-box regulatory element in the upstream region **(Supplemental File 2)**. Owing to the circadian-dependent distinct transition peaks of perireceptor transcripts *i.e*., CYP450 and OBPs (*i.e*., ZT12 for *Anopheles culicifacies* and ZT6 for *Aedes aegypti*) **(Figure 2B, D, E, F),** we hypothesize that the initial physiological events in the peri-receptor space may be more prone towards temporal regulation. Moreover, their distinct dynamic change in diurnal and nocturnal mosquitoes might play key functions in odor detection and rhythmic olfactory sensitivity **(Figure S3D)**.

### Time-of-day has a limited effect on olfactory sensitivity to certain odorants

Next, we asked whether the temporal rhythms of mRNA expression coincided with olfactory sensitivity in both diurnal and nocturnal mosquitoes and compared the temporal Electroantennographic (EAG) sensitivity of nocturnal *An. gambiae* with diurnal *Ae. aegypti* mosquitoes at four different time-ranges *i.e*., ZT1-3, ZT 6-8, ZT12-14, ZT18-20 **(Figure S4A)**. Mosquito antennae were exposed to five doses of 12 different active volatile odorants of floral, human, and cattle origin and representative of terpenoid, phenolic, ketone, aldehyde chemical classes and one chemical blend **(Table S1)**. In the case of *An. gambiae*, the amplitudes of odor-evoked responses were significantly influenced by the doses of all the odorants tested (repeated measure ANOVA, *p* ≤ 2e-16) **(Figure S4B)**. Further analysis was performed for the odorants such as phenol (dose: time, *p* ≤ 0.00229), β-ocimine (dose: time, *p* ≤ 0.0176) and 3-octanol (time, *p* ≤ 0.0441), where time or its interaction with dose were found to be significant **(Supplemental File 2)**. The data indicated that responses to phenol did not vary significantly throughout the day, while responses to β-ocimine and 3-octanol showed significant change as a function of time **(Figure 3A, B)**. *An. gambiae* seems to be most sensitive to 10^-1^ concentration of β-ocimine during ZT6-8 (*p* ≤ 0.0127655) (**Figure 3A**), and 10-2 and 10^-1^ concentration of 3-octanol during ZT1-3 (*p* ≤ 0.0179312, *p* ≤ 0.0204470 respectively) (**Figure 3B**).

**Figure 3:**
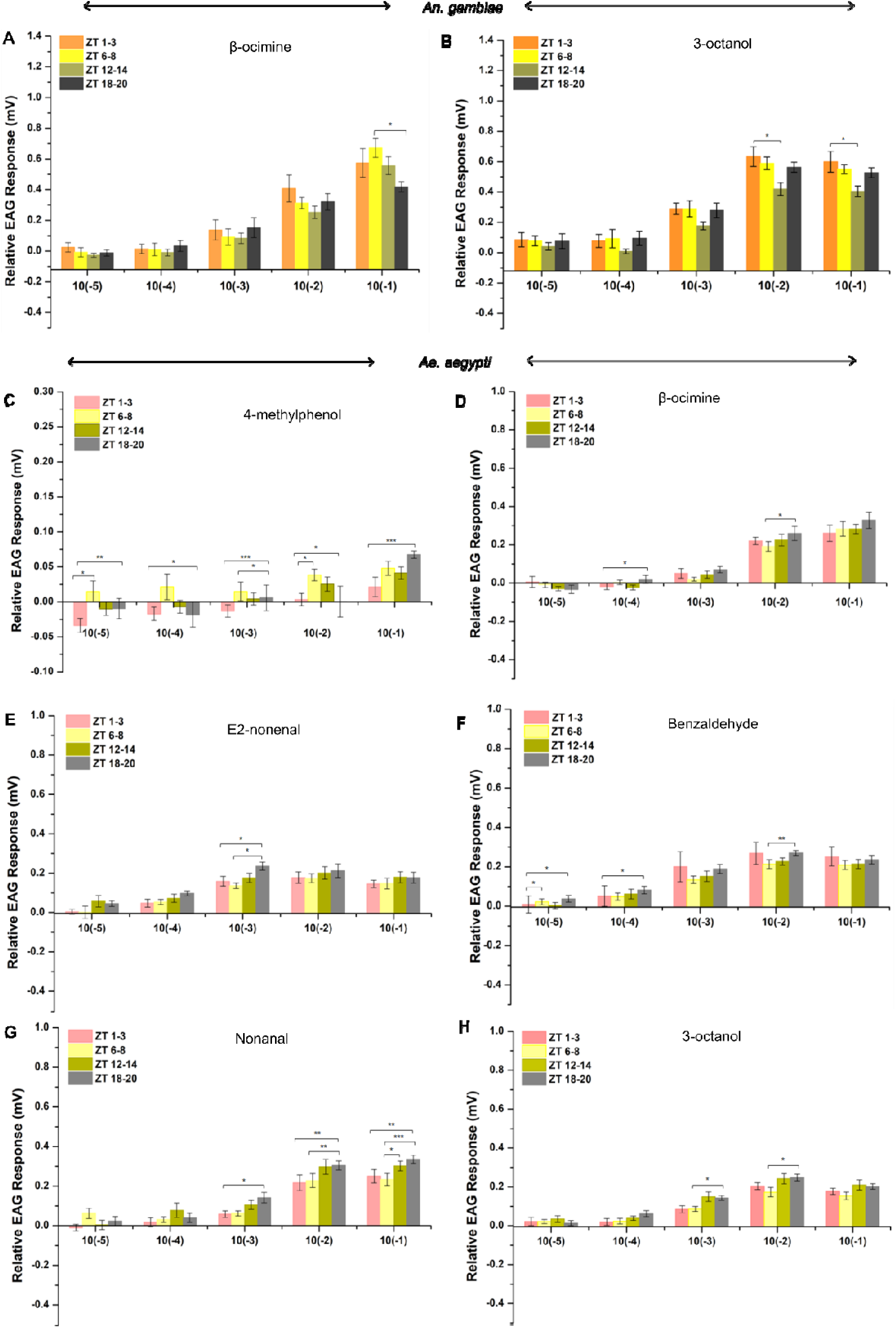
Temporal changes in olfactory sensitivity against different volatile odorants in *An. gambiae* and *Ae. aegypti*. (A) Time-of-day and dose-dependent electroantennographic analysis of adult *An. gambiae* antenna to β-ocimine. (B) Relative EAG responses of *An. gambiae* at different times of day to different stimulus concentrations of 3-octanol. (C-H) Circadian dependent electroantennographic response of adult *Ae. aegypti* antenna against different doses of 4-methylphenol, β-ocimine, E2-nonenal, benzaldehyde, nonanal, and 3-octanol (Different stimulus compounds are indicated on the bottom horizontal axis (10^-5^, 10^-4^, 10^-3^, 10^-2^, 10^-1^). Significant interactions are calculated by one-way ANOVA and Tukey post hoc tests. Significance values are indicated above bar graphs as asterisks (*), where p<=0.05. For all the EAG recordings, N=8-10.

For *Ae. aegypti*, the amplitudes of the odor-evoked responses were also significantly influenced by the doses of all the odorants tested (repeated measure ANOVA, *p* ≤ 2e-16) (Figure S4B). Responses to 4-methyl phenol (*p* ≤ 0.000499), β-ocimine (*p* ≤ 0.0171), E2-nonenal (*p* ≤ 0.0458), benzaldehyde (*p* ≤ 0.0206), nonanal (*p* ≤ 0.00701), sulcatone (dose: time, *p* ≤ 0.0291), 3-octanol (dose: time, *p* ≤ 0.00354) varied significantly as a function of time **(Supplemental File 2)**. *Ae. aegypti* was found to be most sensitive to all the odorants (4-methylphenol, β-ocimine, E2-nonenal, benzaldehyde, nonanal, and 3-octanol) during ZT18-20 except sulcatone (**Figure 3C** **– 3H)**. Interestingly, mosquitoes showed a peak in sensitivity to the 10^-5^ dilution of benzaldehyde during ZT6-8 (*p* ≤ 0.0118817) as well as during ZT18-20 (*p* ≤ 0.0279968) (**Figure 3F**), while for the rest of the odorants, a similar trend of significantly higher response was observed during ZT18-20 **(Figure 3C, 3D, 3E, 3G, 3H)**.

Additionally, the further analysis by considering species identity as the predictor, depicted that odorant responses in both the species were significantly affected by dose and the interaction between species and dose except E2-nonenal (repeated measure ANOVA, *p* ≤ 2e-16) **(Supplemental File 2)**. Among all the odorants, 4-methyl phenol, phenol, β-ocimene, nonanal, and 3-octanol also showed significant variation when species and time interactions were considered. A detailed analysis of those selected odorants revealed that compared to *Ae. aegypti*, *An. gambiae* were most sensitive to (i) 3-octanol (10^-2^, 10^-1^ concentrations) during ZT1-3 and ZT18-20 (**Figure 4A**), (ii) β-ocimine (10^-1^ concentration) during ZT1-3 and ZT6-8 (**Figure 4B**), and (iv) nonanal (10^-1^ concentration) during ZT1-3, ZT6-8 and ZT18-20 (**Figure 4D**). Furthermore, the phenolic compounds (4-methylphenol and phenol) showed significant differences in antennal sensitivity between *An. gambiae* and *Ae. aegypti*. For instance, *An. gambiae* exhibited high sensitivity to 4-methyphenol (10^-5^, 10^-4^, 10^-1^ concentrations) at ZT6-8, ZT12-14, and ZT18-20 (**Figure 4C**), whereas, *Ae. aegypti* displayed enhanced olfactory sensitivity to phenol during the same time points (**Figure 4E**). Additionally, our principal components analysis also illustrates that most loadings of relative EAG responses are higher towards the *Anopheles* observations **(Figure S4C)**. Taken together these data indicate that *An. gambiae* may exhibit higher antennal sensitivity to at least five different odorants tested, as compared to *Ae. aegypti*. Further to check the trend in olfactory responses in another *Anopheline* species, dose-response studies were performed with *An. stephensi*. Similar to *An. gambiae*, a comparatively high amplitude response was also observed in *An. stephensi* **(Figure S4D)**.

**Figure 4:**
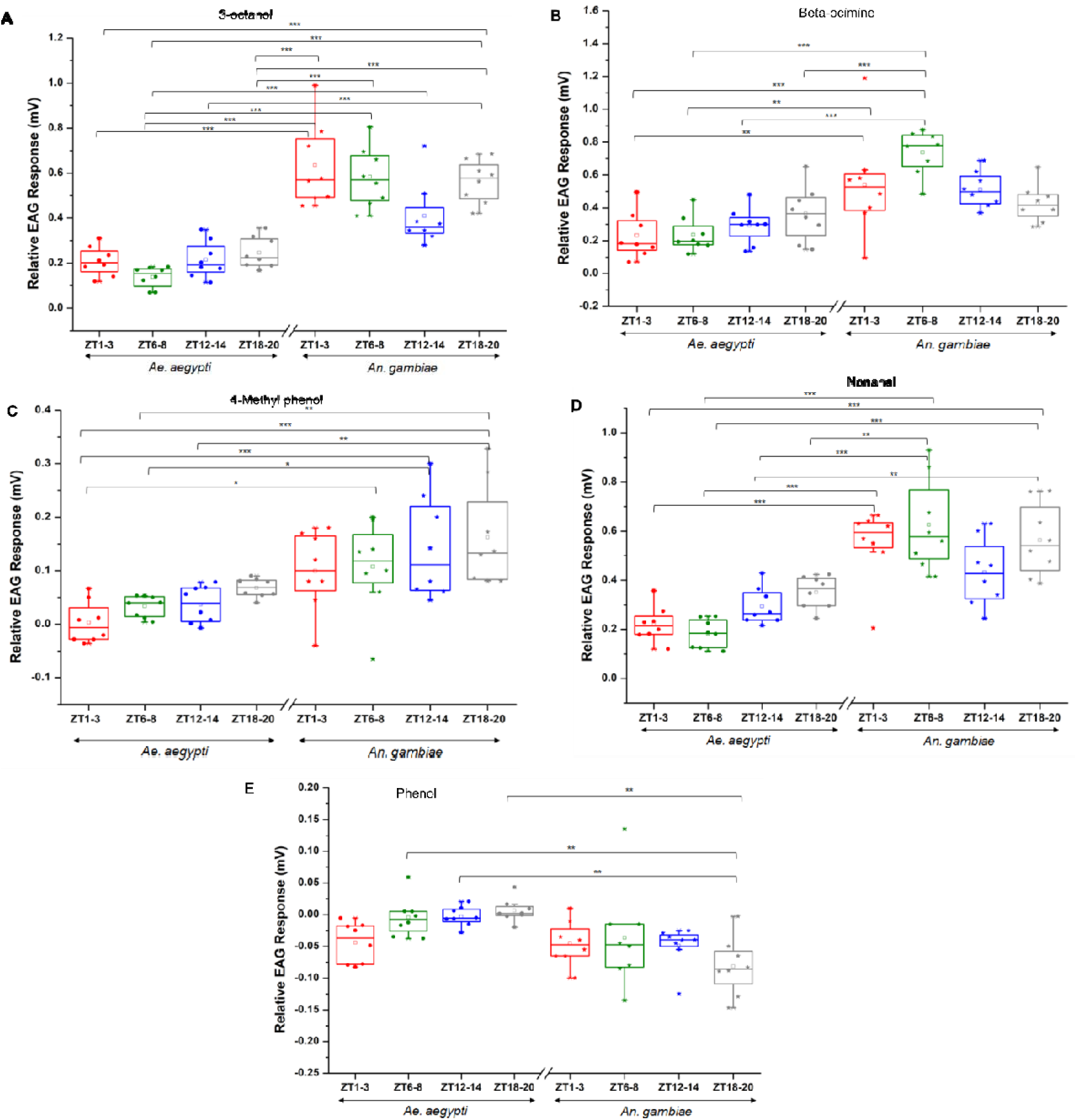
Time-of-day dependent comparison of cross-species antennal sensitivity. (A-E) Circadian-dependent differential antennal sensitivity between *Ae. aegypti* and *An. gambiae* to 3-octanol (10^-2^), β-ocimine (10^-1^), 4-methylphenol (10^-1^), nonanal (10^-1^) and phenol (10^-4^). Significant interactions are derived by two-way ANOVA (between two species differences) followed by post hoc Tukey HSD test between the zeitgeber time points at each concentration of the respective volatile odorant. Significance values are indicated above bar graphs as asterisks (*), where p<=0.05. For all the EAG recordings, N=8-10.

### Knockdown of a cytochrome P450 gene suppresses olfactory sensitivity

Although multiple pathways and several molecular factors are involved in the regulation of olfactory sensitivity in mosquitoes, the time-of-day dependent enrichment of the oxidoreductase category of transcripts, including cytochrome P450 and prophenoloxidase, highlights their possible function in odorant detection. Therefore, functional disruption of cytochrome P450 may influence olfactory sensitivity in the mosquitoes. To test this hypothesis, dsRNA mediated knockdown of one of the significantly upregulated (log_2_2.2-fold) CYP450 genes (AAEL014619), was performed in *Ae. aegypti* **(Figure 5A, B)**. Interestingly, a notable decrease in relative EAG response against all the six odorants tested was observed **(Figure 5C, S5A)**, indicating the involvement of the CYP450 in odorant detection. To further establish this proof-of-concept in *An. gambiae*, three potent CYP450 inhibitors, aminobenzotriazole(*52*), piperonyl butoxide(*53*), and schinandrin A (*54*), was applied topically on the head capsule of 5-6-day-old female mosquitoes (**Figure 5D**). A significant decrease in the EAG amplitude was observed in aminobenzotriazole and piperonyl butoxide treated mosquitoes for all six odorants tested (**Figure 5E** **& Figure S5C)**. Schinandrin A also induced a significant decrease in olfactory sensitivity but had lower potency than the other two inhibitors **(Figure S5B)**. Next, we carried out similar inhibitor studies in *Ae. aegypti* mosquitoes. Though inhibitor application significantly affected odorant sensitivity in *Ae. aegypti,* a large inter-individual variation in the response was observed (**Figure 5F**). Taken together, inhibitor treatment and RNAi-mediated knockdown of CYP450 proteins provide compelling evidence supporting the crucial role of CYPP450 in the odor detection process in mosquitoes.

**Figure 5:**
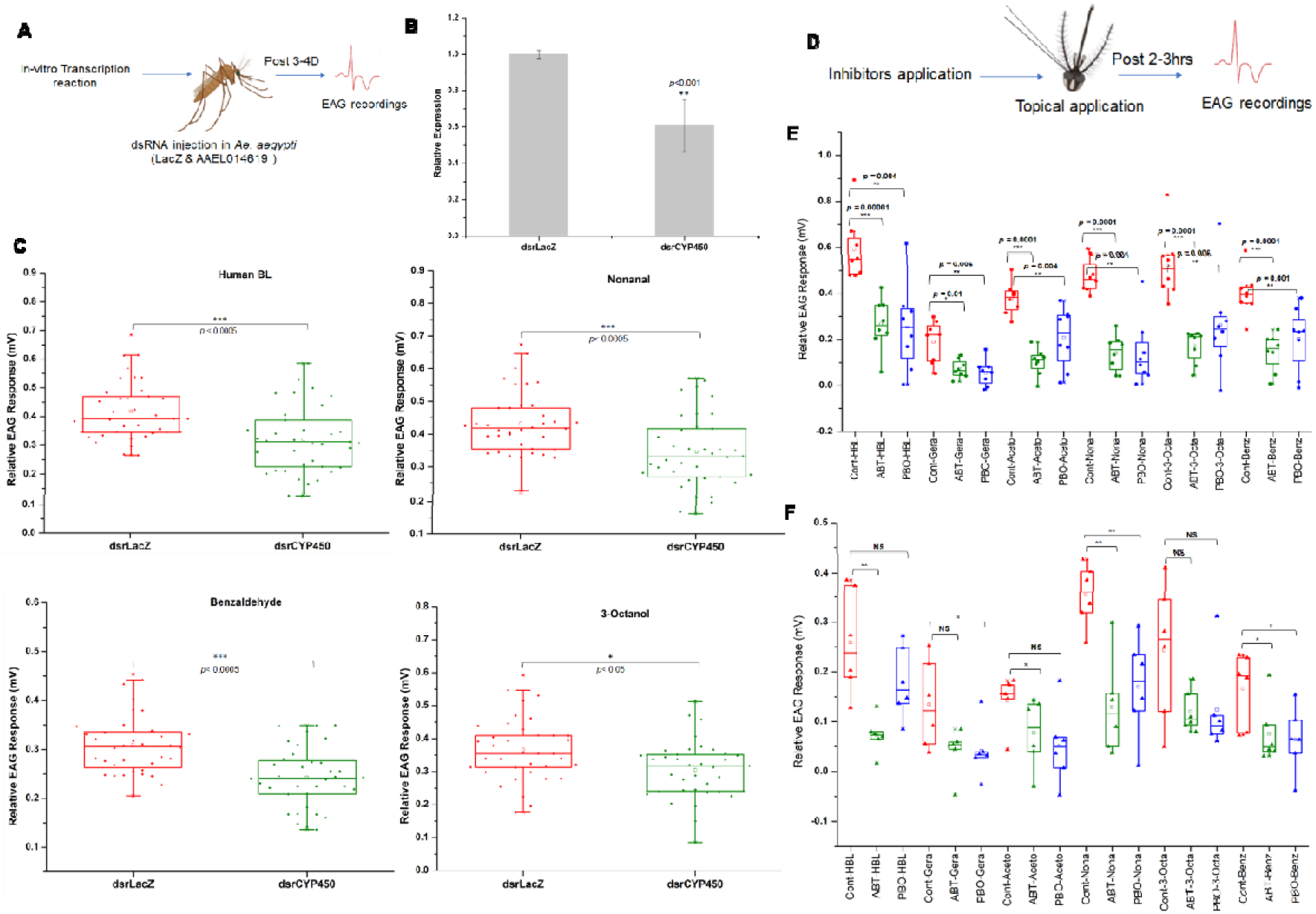
Suppression of cytochrome P450 (CYP450) function reduces the olfactory sensitivity in mosquitoes. (A) Schematic representation of RNAi-mediated knock-down and functional EAG assays. (B) Real-time PCR-based validation of CYP450 (AAEL014619) silencing in *Ae. aegypti*. Decapitated heads of control and knock-down mosquitoes were collected for RT-PCR validation following EAG recordings. A total of 3-5 heads (n), that showed change in EAG response, were pooled in a single tube to look for knock-down efficiency. The number of biological replicates, N=4 for RT-PCR experiment. Statistically significant value was calculated by students’ t-test. (C) Relative EAG response of dsrLacZ and dsrCYP450 treated mosquitoes against four different odorants. The odor response was tested with 40 individual mosquitoes of each group (N=40). (D) Schematic representation of inhibitor studies in mosquitoes. (E) Relative EAG response of naive (70% EtOH treated) and inhibitors treated *Ae. aegypti* (N=6) against six different odorants. ABT: Aminobenzotriazole, PBO: Piperonyl butoxide. Volatile odorants: HBL-human blend, Gera-geraniol, Aceto-acetophenone, Nona-nonanal, 3-Octa-3octanol, Benz-benzaldehyde. (F) Comparative EAG response of naive (70% EtOH treated) and inhibitors treated *An. gambiae* (N=6) against six different odorants. ABT: Aminobenzotriazole, PBO: Piperonyl butoxide. Volatile odorants: HBL-human blend, Gera-geraniol, Aceto-acetophenone, Nona-nonanal, 3-Octa-3octanol, Benz-benzaldehyde. Statistical significance was tested by Mann Whitney test for individual odorants and p-values are mentioned in the plot.

### Daily temporal modulation of neuronal serine protease impacts mosquito’s olfactory sensitivity

Both external stimuli and internal circadian inputs have a significant impact on the oscillation of synaptic structure and function(*32*), but, how circadian plasticity remodels brain functions in diurnal and nocturnal mosquitoes has not yet been explored. Therefore, to gain an insight into the molecular mechanism of circadian-guided synaptic plasticity, we performed a functional annotation of the brain tissue transcriptome data of both diurnal and nocturnal mosquitoes. A gene-set enrichment analysis of the upregulated genes revealed an over-representation of the functional categories of UDP-glucosyltransferase activity, small molecule metabolic process, oxidoreductase, and monooxygenase activity during ZT6-ZT12, as well as extracellular region and extracellular matrix during ZT6 (**Figure 6A**) in the brain of *An. culicifacies*. Similar to the peripheral olfactory system, the pathway prediction analysis showed a notable enrichment of the xenobiotic/drug metabolism pathway, the steroid hormone biosynthetic pathways, pentose, and glucuronate interconversions, as well as the ascorbate and aldarate metabolism pathway during ZT12. **(Fig. S6A)**. Detailed functional annotation analysis of the oxidoreductase family suggested the predominant expression of CYP450 and prophenoloxidase genes during ZT12 **(Supplementary File 1)**. While further studies are needed to comprehend how time-of-day dependent enhancement of CYP450 influences temporal regulation of steroid and xenobiotic metabolism in the brain of mosquitoes, the previous observations of topical application of inhibitors do not preclude the chances of the inhibitor absorption into the brain. Therefore, the observed phenotype of a significant decrease in olfactory responses following inhibitor treatment may be a cumulative systemic effect of inhibition of both brain and peripheral olfactory CYP450 genes.

**Figure 6:**
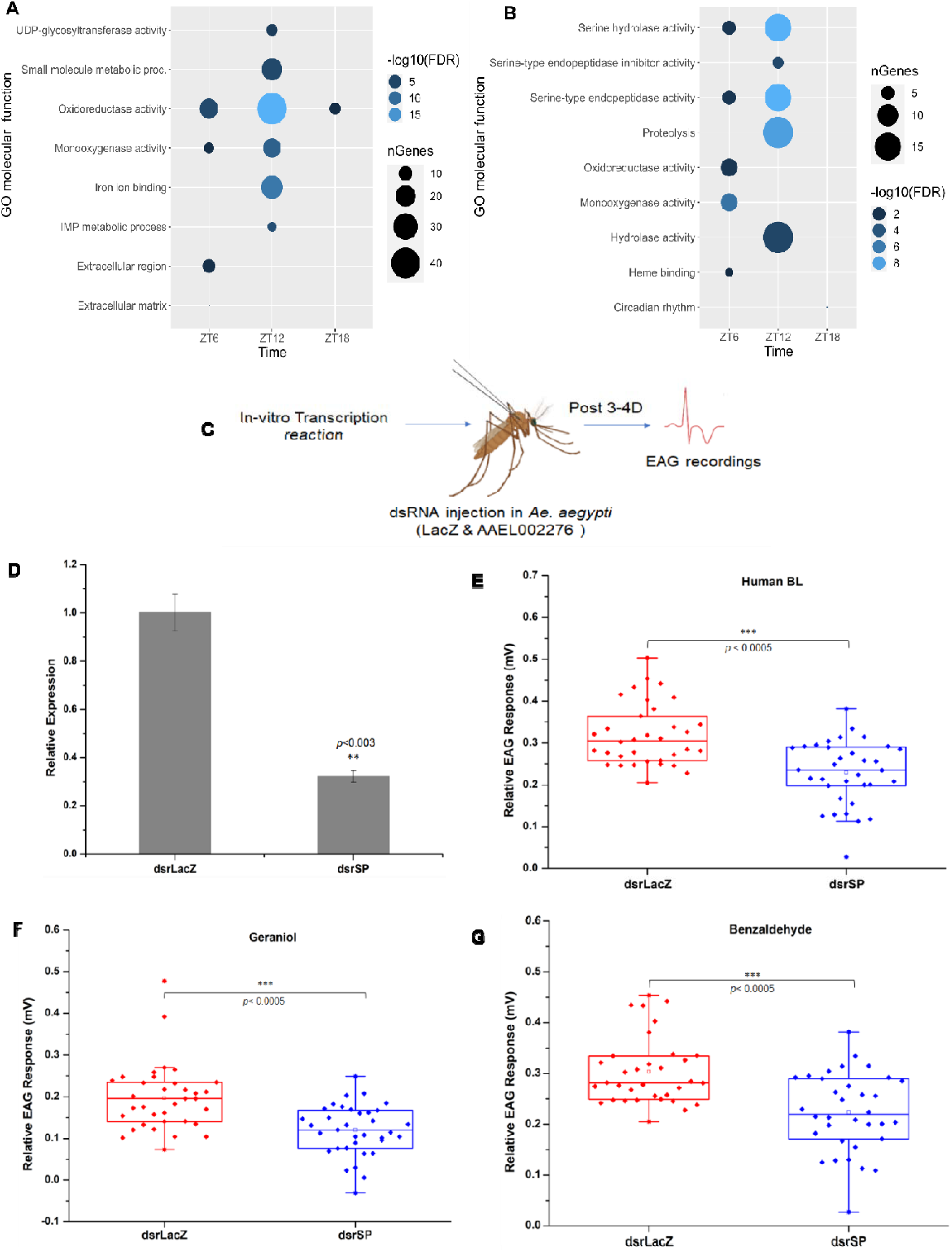
Gene ontology and functional annotation analysis of brain transcriptome of nocturnal and diurnal mosquitoes. (A-B) Gene ontology and functional annotation analysis of differentially upregulated genes in *An. culicifacies* (A) and *Ae. aegypti* (B). The green colored box highlights the most abundantly upregulated genes. (C) Schematic representation of RNAi mediated knockdown in *Ae. aegypti*. (D) RT-PCR validation of successful knock-down of the serine protease (SP) gene (AAEL002276) in *Ae. aegypti*. A total of 3-5 heads (n), that showed change in EAG response, were pooled in a single tube to assess knock-down efficiency. The number of biological replicates, N=4, for RT-PCR experiment. Statistically significant value was calculated by students’ t-test. (E-G) Relative EAG response of naive (dsrLacZ injected) vs knock-down (dsrSP) *Ae. aegypti* against human blend (E), geraniol (F), benzaldehyde (G). The odor response was tested with 40 individual mosquitoes of each group (N=40). Statistical significance was tested by Mann Whitney test for individual odorants and p-values are mentioned in the plot.

Though, differential gene expression analysis exhibited a limited number of differentially expressed genes in the brain of *Aedes aegypti* **(Fig. S2B)**, serine-type endopeptidase and hydrolase activity categories of genes were found to be highly enriched at ZT12, and the functional categories of oxidoreductase and monooxygenase activity are overrepresented at ZT6 (**Figure 6B**). Similar to *An. culicifacies*, CYP450, and propenoloxidase genes predominantly occupy the monooxygenase category (**Supplementary File 1)**. Notable enrichment of CLIP-domain serine protease and serine protease inhibitor genes was observed under the ontology category of serine-type endopeptidase activity **(Figure 6B; Figure S6B)**. Although the function of serine protease and serine protease inhibitors in mosquitoes have mostly been studied with respect to their role in innate immune response (*55*, *56*), their physiological relevance is underexplored. It has become increasingly evident that these groups of proteins are widely expressed in brain cells and have a long-term impact on the plasticity, memory, cognition, and behavior of the organism (*39*, *45*, *57*). Therefore, to study the effect of neuronal serine protease on circadian-dependent olfactory responses, we carried out RNAi-mediated knock-down of one of the serine-protease (SP) genes (AAEL002276) in *Ae. aegypti* (that showed log_2_2.2-fold upregulation during ZT12), and assessed changes in olfactory sensitivity by EAG (**Figure 6C**). Notably, the SP-knock-down mosquitoes displayed reduced antennal response when compared with the dsrLacZ-treated mosquitoes (**Figure 6D**) against the odor blend and all six odorants tested (**Figure 6E-G** **and Figure S6C-E)**. Our finding highlights that daily temporal modulation of neuronal serine-protease may have important functions in the maintenance of brain homeostasis and olfactory odor responses.

## Discussion

The evolution of the species-specific specialized neuro-olfactory system and its distinct temporal regulation in mosquitoes facilitates their coexistence in the same ecological niche by minimizing inter-species competition (*15*, *46*, *47*). However, the underlying molecular mechanism that synchronizes time-of-day dependent regulation of olfaction in mosquitoes is poorly understood. Since diurnality and nocturnality are the two contrasting behavioral patterns of *Aedes* and *Anopheles* mosquitoes, we aimed to decode the differential molecular rhythms of their neuro-olfactory system.

### Circadian oscillation of peri-receptor proteins regulates olfactory sensitivity

In this study, we provide initial evidence that the daily rhythmic change in the olfactory sensitivity of mosquitoes is tuned with the temporal modulation of molecular factors involved in the initial biochemical process of odor detection *i.e.,* peri-receptor events. The findings of circadian-dependent elevation of xenobiotic metabolizing enzymes in the olfactory system of both *Ae. aegypti* and *An. culicifacies* are consistent with previous literature (*26*, *31*), and we postulate that these proteins may contribute to the regulation of odorant detection in mosquitoes. Given the potentially larger odor space in mosquitoes (like other hematophagous insects) (*16*, *58*), it can be hypothesized that detection of any specific signal in such a noisy environment, mosquitoes may have evolved a sophisticated mechanism for rapid (i) odor mobilization and (ii) odorant clearance, to prevent anosmia (*24*). We noticed peak transcriptional changes of the peri-receptor proteins (ZT12 in *An. culicifacies* and ZT6 in *Ae. aegypti*), specifically XMEs, during the preceding phase of active olfaction/navigation (ZT16-ZT-20 in *An. culicifacies* and ZT10-12 in *Ae. aegypti*) (*50*, *51*). Previous microarray analysis by Rund *et.al*. also demonstrated circadian-dependent enrichment of several detoxification categories of genes in the heads of *An. gambiae* (*26*). Additionally, a complete cease in the OBPs expression was also observed during early morning (ZT0) and mid-night (ZT18) in *An. culicifacies* and *Ae. aegypti*, respectively. Taken together, we hypothesize that circadian-dependent activation of the peri-receptor events may modulate olfactory sensitivity and are key for the onset of peak navigation time in each mosquito species.

To correlate the molecular rhythm and modulation of olfactory sensitivity in mosquitoes, we assessed the temporal rhythms of odor responses through electroantennography. We conducted a dose-dependent analysis to trace the interaction between doses and temporal sensitivity between a nocturnal and diurnal species. Due to technical limitations, and considering the substantial data on the circadian-dependent molecular rhythmicity (*25*, *26*, *59*), the electroantennography study was performed on *An. gambiae* G3 as a representative of nocturnal species instead of *An. culicifacies*. Accounting for all the cross-comparison among species, time-of-day, and concentration of the odorants, a noteworthy interaction between species and doses was identified. In contrast to *An. gambiae*, the time-dose interactions had a higher significant impact on the antennal sensitivity of *Ae. aegypti*. *An. gambiae* showed a conserved pattern in the daily rhythm of olfactory sensitivity, peaking at ZT1-3 and ZT18-20. This observation does not fully harmonize with previous studies in *An. gambiae*, in which the highest antennal sensitivity was reported at ZT16 (*25*), overlapping with the highest expression of olfactory proteins. Together these data, we interpret that mosquito’s olfactory sensitivity possibly does not follow a fixed temporal trait and is susceptible to change depending on (a) the presence of different chemical odorants and mosquitoes need to detect those to drive various behavioral repertoires (*60*), (b) diverse ecological/environmental habitats (*61*, *62*), (c) host-availability and (d) presence of xenobiotics (*63*, *64*). Moreover, we hypothesize that under standard insectary conditions, mosquitoes may not need to exhibit foraging flight activity either for nectar or blood, and during the time course, it may minimize their olfactory rhythm, which is obligately required for wild mosquitoes. Previous literature also indicated that depending on the task demands foragers and nurses’ ants showed plasticity in their behavioral rhythms which also varied between an isolated and social individual (*65*). Therefore, it is reasonable to think that the mosquitoes used for EAG studies may have adapted well under insectary settings and, hence carry weak olfactory rhythm.

*Aedes aegypti* displayed a peak in antennal sensitivity at ZT18-20 to the higher concentrations of plant and vertebrate host-associated odorants tested. Given the time-of-day dependent multiple peaks (at ZT6-8 and ZT18-20 for benzaldehyde and at ZT12-14 and ZT18-20 for nonanal) in antennal sensitivity to different odorants, our data supports the previous observation of bimodal activity pattern of *Ae. aegypti* (*50*). We also observed that, compared to the chemical blend used in the current study, the individual chemical odorants showed more significant temporal rhythmicity. Although the overall relative EAG responses to 4-methylphenol were much lower as compared to other odorants, mosquitoes displayed higher antennal sensitivity to lower concentrations of 4-methylphenol (10^-5^, 10^-4^, 10^-3^, 10^-2^) at ZT6-8. Previous reports indicate that 4-methylphenol is a broad-spectrum odorant having function in host-seeking, blood-feeding, and oviposition site selection (*16*, *66*, *67*). Thus, it is plausible that the antennal response in *Ae. aegypti* is very sensitive to lower concentrations of 4-methylphenol at ZT6-8, facilitating their diurnal activity. Interestingly, our species-time interaction studies revealed that *An. gambiae* exhibits time-of-day dependent significantly high antennal sensitivity to at least four chemical odorants compared to *Ae. aegypti*, except phenol. Similar observations were also noticed with *An. stephensi.* Although additional work with diverse chemical odorants and other mosquitoes is needed to comprehend the species-specific difference in antennal sensitivity, our preliminary data indicate that *Anopheles* spp. may possess comparatively higher olfactory sensitivity to a substantial number of odorants as compared to *Aedes* spp.

Furthermore, corroborating with previous studies, we also noticed a higher abundance of rhythmic genes including the oxidoreductase category of genes were enriched in *Anopheles* species compared to *Aedes* (*26*). Whether this higher abundance of detoxification enzymes in the olfactory sensilla has any impact on high antennal sensitivity, remains to be explored. To validate the proof-of-concept, we combined RNAi and electroantennographic approach to measure the olfactory sensitivity of both *Anopheles gambiae* and *Aedes aegypti*. A significant decrease in odorant sensitivity for all the volatile odors tested in the CYP450-silenced *Ae. aegypti*, highlighted the possible role of CYP450s in odorant detection. However, the compensatory actions of other CYP450 homologs in olfaction cannot be ignored. Thus, to get more informative data, we applied potent CYP450 protein inhibitors on the head capsule of mosquitoes including the antennae. As these inhibitors, upon absorption through the antennal cuticle, block all the CYP450 expressed in the sensilla, we anticipated a more significant effect on olfactory sensitivity. As expected, a drastic reduction of olfactory sensitivity against all tested odorants was recorded with two potent inhibitors, aminobenzotriazole and piperonyl butoxide, in both *An. gambiae* and *Ae. aegypti*. Taken together, our data demonstrate that CYP450 may have a crucial role in odorant detection, which influences time-of-day-dependent olfactory sensitivity in mosquitoes.

### Neuronal serine protease consolidates brain function and olfactory detection

Despite their tiny brain size, mosquitoes, like other insects, have an incredible power to process and memorize circadian-guided olfactory information (*7*). While the morphological structure of neurons, peri-synaptic space, and glial cells are plastic and prone to change during exposure to internal and external stimuli (*35*, *48*, *49*), circadian inputs may also alter pre-and post-synaptic morphology, as demonstrated in *Drosophila* (*32*, *48*). Furthermore, it is becoming evident in flies and mammals that neuronal protease (*e.g.,* serine proteases, matrix-metalloproteases) mediated remodeling of extracellular matrix molecules (ECM) at the synapse is important for the homeostatic process of the brain, which facilitate new synaptic connections and has potential implication in synaptic signaling (*38*, *39*). Induced expression of serine protease, tissue plasminogen activator (tPA), during the motor learning process, was found to be responsible for long-term potentiation (LTP) in mice (*39*, *68*). Furthermore, a number of studies implicated that the collaborative actions of neuronal serine protease and serine protease inhibitors (serpins) can directly influence the activation of a class of G-protein coupled receptors called protease-activated receptors (PAR) in the nervous system (*45*, *69*). Proteolytic activation of PAR proteins modulates synaptic efficacy and plasticity by regulating glutamatergic and GABAergic neuro-transmission (*45*, *69*). However, the functional evaluation of neuronal serine protease in invertebrates is only limited to the fruit fly *Drosophila melanogaster*, where it was found to modulate blood-brain barrier structure by sensing the nutritional status (*57*).

Circadian-guided natural upregulation of several serine proteases and their inhibitor proteins in the brain of *Aedes* mosquitoes during the dusk transition phase prompts us to hypothesize its function in the circadian regulation of synaptic plasticity and odor detection in mosquitoes. Though, it is hard to comment on the exact mechanism of actions (either through ECM remodeling or PAR-mediated activation or both) of these serine proteases in mosquito brain, suppression of olfactory sensitivity in serine protease knock-down mosquitoes indirectly indicates its possible role in synaptic signal transmission and plasticity. Furthermore, accounting for the “State-Clock Model”, the temporal oscillation of the sleep-wake cycle of any organism is managed by the encoding of experience during wake, and consolidation of synaptic change during inactive (sleep) phases, respectively (*70*). We speculate that after the commencement of the active phase (ZT6-ZT12), the serine peptidase family of proteins in the brain of *Ae. aegypti* mosquitoes may play an important function in consolidating brain actions (after ZT12) and aid circadian-dependent memory formation. In *Anopheles* mosquitoes, the gene ontology analysis does not show the enrichment of serine peptidase family proteins, however detailed functional annotation analysis revealed multifold (>log_2_3.1-fold) enhancement of a few proteases, including serine protease during the early morning (ZT0). As *An. culicifacies* exhibit their peak activity during the night (∼ZT16-ZT20), they probably consolidate their brain function during the dawn-transition phase. Additionally, a rhythmic secretion of steroid hormones and release of neuromodulators were reported to modulate neuronal excitability and plasticity(*70*, *71*). A notable enrichment of oxidoreductase family proteins, including CYP450, in the brain of both diurnal and nocturnal mosquitoes indicates the possible role of these proteins in neuro-hormonal rhythmicity. However, elucidating the mechanism of neuro-hormone-guided change in neurotransmission requires future efforts.

### Limitation of the study

In this study, we have discussed/demonstrated an association between the abundance of genes encoding for perireceptor proteins and olfactory sensitivity. However, (1) our results do not allow us to conclude that XMEs play a central role in the regulation of olfactory rhythm and odor detection. Demonstrating this correlation requires further behavioral studies on mosquitoes and *in vitro* biochemical characterization of the CYP450 proteins. Furthermore, due to technical limitations, we used different *Anopheles* species for our electroantennography studies. Thus, we cannot rule out the possibility of species-specific distinct temporal olfactory sensitivity of mosquitoes (*An. culicifacies* vs *An. gambiae*). Furthermore, (2) we exclusively focused on EAG studies to correlate the function of serine protease in synaptic functions and plasticity of mosquitoes. Subsequent work on electrophysiological and neuro-imaging studies are needed to demonstrate the role of neuronal-serine proteases in the reorganization of perisynaptic structure.

### Closing remarks

Targeting multiple disease vectors together at a particular time of the day may reduce the efficacy of the vector control procedure. Decoding the molecular mechanism of plasticity in daily rhythmic behaviors of different mosquito species may have important implications in vector management programs, since targeting all vectors together, only at a particular time of the day, may reduce the efficacy of the treatment procedure. Our work establishes the important role of peri-receptor proteins (OBPs and XMEs) in determining the olfactory sensitivity in both diurnal and nocturnal mosquitoes. Moreover, we propose that neuronal serine protease may have a crucial function in consolidating brain actions that in consequence influence synaptic plasticity and olfactory sensitivity. In summary, our study advances the conceptual understanding of odor detection mechanisms in mosquitoes. Therefore, detailed knowledge about the distinct peak-transition phase of XMEs and other neuro-olfactory factors in different mosquito species could be helpful for the rational designing of intervention strategies.

## Material and Methods

### Mosquito rearing and maintenance

*Anopheles culicifacies*, sibling species A, were maintained at 80% relative humidity and 27 ± 2 °C under a 12 h/12 h light: dark cycle, with a 1 h dawn and a 1 h dusk transition phase, in the central insectary facility of the ICMR-National Institute of Malaria Research (ICMR-NIMR). For colony maintenance, 10% sucrose solution was provided *ad libitum*, and adult female mosquitoes were blood-fed on a live rabbit at night. All protocols for rearing and maintenance of the mosquito culture were approved by the ethical committee of the ICMR-NIMR(*72*). For *Ae. aegypti*, the laboratory-adapted strain of wild-caught mosquitoes from Raipur of Chhattisgarh state was used, and maintained under similar insectary conditions as *An. culicifacies,* albeit blood-fed during daytime. For EAG studies, *An. gambiae* (G3), *An. stephensi* and *Ae. aegypti* (Rockefeller) were used and reared under similar insectary conditions at the Swedish University of Agricultural Sciences.

### Tissue collection, RNA isolation, and transcriptome sequencing analysis

To understand the molecular rhythm of the olfactory and the brain system, complete olfactory tissue, including antennae, maxillary palps, proboscis, and labium, as well as the brain were dissected from 5-to-6-days old adult female mosquitoes (same cohort) at four different zeitgeber times, ZT0, ZT6, ZT12 and ZT18. At first, cold anesthetized mosquitoes were decapitated, and then the olfactory and brain tissues were dissected and pooled from 20-25 mosquitoes in Trizol(*73*) (RNAiso Plus, Takara Bio, Kusatsu, Japan) for a total of three biological replicates. Total RNA was extracted from the tissues, and cDNA libraries were prepared using NEBNext® UltraII RNA library preparation kit (NEB, New England Biolabs, MA, USA) using the manufacturer’s protocol. For multiplexing of the samples, NEBNext® Multiplex Oligos for Illumina® (NEB, New England Biolabs) were used. Sequencing was performed using Illumina NextSeq550 platform. Next, the raw reads were demultiplexed and converted to FASTQ format using bcl2fastq conversion software (https://support.illumina.com/downloads/bcl2fastq-conversion-software-v2-20.html). After quality check and adaptor trimming, the sample-specific reads were aligned to the reference genome of *An. culicifacies* (VectorBase-54-AculicifaciesA-37-Genome; https://www.vectorbase.org/) and *Ae. aegypti* (VectorBase-54-AaegyptiLVP-AGWG-Genome; https://www.vectorbase.org/), respectively, by using STAR-alignment tool(*74*). The transcriptome data of *An. culicifacies* showed 70-80% alignment rate, whereas the *Ae. aegypti* data showed ∼90% alignment. Finally, the differential gene expression analysis was performed using the DESeq2 package of R version 4.2(*75*). For functional annotation and gene enrichment analysis, ShinyGO 0.741 tool was used(*76*). Heatmaps were prepared by using either an online available heatmapper tool (http://www.heatmapper.ca/) or pheatmap package in R version 4.2 (*77*).

### Odor preparation for electroantennographic recordings

For the EAG analysis, volatile compounds of different origin (plant, vertebrate or both) and chemical class (monoterpenoids, aromatics, aldehydes, and ketones) were selected. Neat volatile chemicals, including (+)-camphor (CAS: 76-22-2; 96%), geraniol (CAS: 106-24-1; 98%), 4-methyl phenol (CAS: 106-44-5; 99%), acetophenone (CAS: 98-86-2; 99%), phenol (CAS: 108-95-2, 99%), β-ocimene (CAS: 13877-91-3; 90%), *E*2-nonenal (CAS: 18829-56-6, 97%), benzaldehyde (CAS: 100-52-7, 99%), nonanal (CAS: 124-19-6; 95%), limonene (CAS: 138-86-3; 97%), sulcatone (CAS: 110-93-0; 99%), and 3-octanol (CAS: 589-98-0; >95%) were obtained from Sigma-Aldrich (Stockholm, Sweden) at the highest purity available, and dilutions were prepared in redistilled hexane (>95%, Sigma-Aldrich), except for camphor and phenol for which double distilled water was used. The previously tested and validated synthetic human blend that mimics the ratio and composition of the compounds of the human headspace, was used(*27*). Five different dilutions, from 10^−1^ to 10^−5^, of each odorant and odor were prepared by serial dilution and stored at -4 °C. An aliquot of 10 µl of each dilution was loaded onto a piece of Whatman filter paper (5 mm x 15 mm) inserted into a glass Pasteur pipette, and the solvent was allowed to evaporate for at least 15 min under a fume-hood before use. To avoid the depletion of the odorant molecules, each odor cartridge was used for four individual recordings with an incubation period of 30 min in between.

### Electroantennographic recordings

For EAG recordings, 5-to-6-days old non-blood-fed adult female mosquitoes of the same cohort were used. Four different ranges of zeitgeber times (ZT1-3, ZT6-8, ZT12-14, ZT18-20) were selected for the recordings, corresponding closely to those used in the RNA-Seq analysis. The head of the mosquito and the distal tip of one antenna were cut off. Then, the tip of the reference electrode, filled with Beadle-Ephrussi ringer solution, was inserted into the foramen of the head. The distal end of the antenna was inserted into a Beadle-Ephrussi ringer-filled recording electrode, connected to a high-impedance probe (10x, Ockenfels Syntech, Buchenbach, Germany) using a micromanipulator. The antennal preparation was placed in a continuous humidified and charcoal filtered air stream (1 l/min), delivered via a glass tube ending 0.5 cm from the antennal preparation. The recording signals were fed into an IDAC2-USB box (Ockenfels Syntech) and then visualized and stored on a PC(*78*). The antennal responses (in mV) of all the recordings were manually analyzed using the GC-EAD 2011 software (Ockenfels Syntech). The antennal preparation was exposed to the increasing doses of the five dilutions of the different odorants and odor that were puffed at 15-20 sec intervals, with a solvent control in between each stimulus. Mosquitoes that were recorded at ZT12-14 and ZT18-20, were kept in the dark, and the EAG recordings were performed under dark conditions. The normalized antennal responses, that were calculated by subtracting the average solvent control response, are considered for further analysis.

### Data analysis

To assess significant changes in olfactory sensitivity, to ecologically relevant volatile odorants, in response to the temporal rhythm, a repeated measures ANOVA for each volatile odorant was carried out using dose and time as predictors in the model, in which individual identity of the mosquito (ID) was nested with dose for the two-mosquito species. To conduct a comparative analysis between the two species for every volatile odorant, a similar analysis was carried out as above while adding species identity as an additional predictor. If the time of day or its interaction terms were found to influence the antennal sensitivity a one-way ANOVA (within single species differences) or two-way ANOVA (between two species differences) followed by post hoc Tukey HSD test between the zeitgeber time points at each concentration of the volatile odorant (dose) was performed. To check the normality of the datasets, Shapiro Wilks test was carried out, or data were square root transformed to correct for non-normal distributions. All the mentioned analyses were conducted using the software R (version 4.3.1).

### dsRNA-Mediated gene knock-down

To suppress the expression of cytochrome P450 (AAEL014619) and serine protease (AAEL002276) genes, an *in vitro* transcription reaction was performed to prepare dsRNA by using Megascript RNAi kit (Ambion by life technologies; Texus, USA). Purified dsRNA was injected into cold anesthetized 1–2-day(s)-old *Ae. aegypti*. For the control group, age-matched mosquitoes were injected with LacZ dsRNA (bacterial origin). Three-to-four days post-dsRNA injection, both groups of mosquitoes were used for EAG recordings. Individual heads of the control and knock-down mosquitoes, which were used for recording, were collected in Trizol (RNAiso Plus, Takara Bio, Kusatsu, Japan) for RNA extraction and cDNA preparation. The silencing efficiency was evaluated through quantitative real-time PCR (CFX96 real-time thermocycler; BioRad; Berkeley, USA). A standard PCR program of 95 °C for 5 min, 40 cycles of 95 °C for 10 s, 60 °C for 15 s and 72 °C for 22 s, followed by a melt curve analysis in 0.5 °C increments from 55 °C to 95 °C for 15 s each. For functional validation, EAG recordings were performed from 40 mosquitoes of the control and treated groups. To assess a change in olfactory sensitivity, the data was analyzed using the Mann-Whitney test.

### Inhibitor treatment

In order to suppress the function of all CYP450 expressed in the olfactory system, three potent inhibitors, aminobenzotriazole (Sigma-Aldrich; Cat. No. - A3940-50MG)(*52*), piperonyl-butoxide (Sigma-Aldrich; Cat. No. - 78447-50MG)(*53*) and schinandrin A (Sigma-Aldrich; Cat. No.SMB00368-10MG)(*54*) were applied topically on the head capsule of adult 5-6-day old *An. gambiae* and *Ae. aegypti*. At first, the inhibitors were dissolved in absolute ethanol at a concentration of 20 mg/ml and then half-diluted in phosphate-buffer-saline. A 50 nl working dilution of the inhibitors were applied on the head capsule of the mosquitoes by using a nano-injector (Drummond Scientific Company, Pennsylvania, USA). After inhibitor application, the mosquitoes were kept in the insectary for 2-3 h. For the control group, 70% ethanol was applied on the head capsule of mosquitoes collected from the same cohort, and incubated for the same time period. EAG recordings were performed with the inhibitor and control mosquitoes against six different odorants. No mortality or alteration in flight activity was observed following application of either the inhibitors or the ethanol.

## Data access

The RNA-Seq data and the analyzed differential gene expression data are deposited the Gene Expression Omnibus (GEO) database under accession no. GSE238168.

## Ethics Statement

All protocols for rearing and maintenance of the mosquito culture were approved by the ethical committee of the institute, ICMR-National Institute of Malaria Research(*72*).

## Acknowledgments

We are thankful to the NGS facility, IISER Pune for sequencing support. TD was financially supported by SERB-National Post-Doctoral Fellowship grant, Department of Science and Technology (Grant No. PDF/2020/000887). We are thankful to EMBO-Scientific-Exchange Grant (Grant No. 9563) for travel and financial support for visiting SLU, Sweden. We are also thankful to Anjani Rao for library preparation and sequencing support and Adam Flöhr, SLU, for statistical analysis.

## Authors’ contributions

Conceived and designed the experiments: TDD, JP, RI, KK; Performed the experiments: TDD, JP; Analysed the data: TD, JP, SG, RI, KK; Contributed reagents, materials, analysis tools: RD, KMP, OPS, RI, KK; Wrote the paper: TD, KK, JP, RI, SG, RD. Revised the manuscript: All authors read and approved the final version of the manuscript.

