## Supplementary Information for "Circadian oscillation of perireceptor events influence olfactory sensitivity in diurnal and nocturnal mosquitoes"

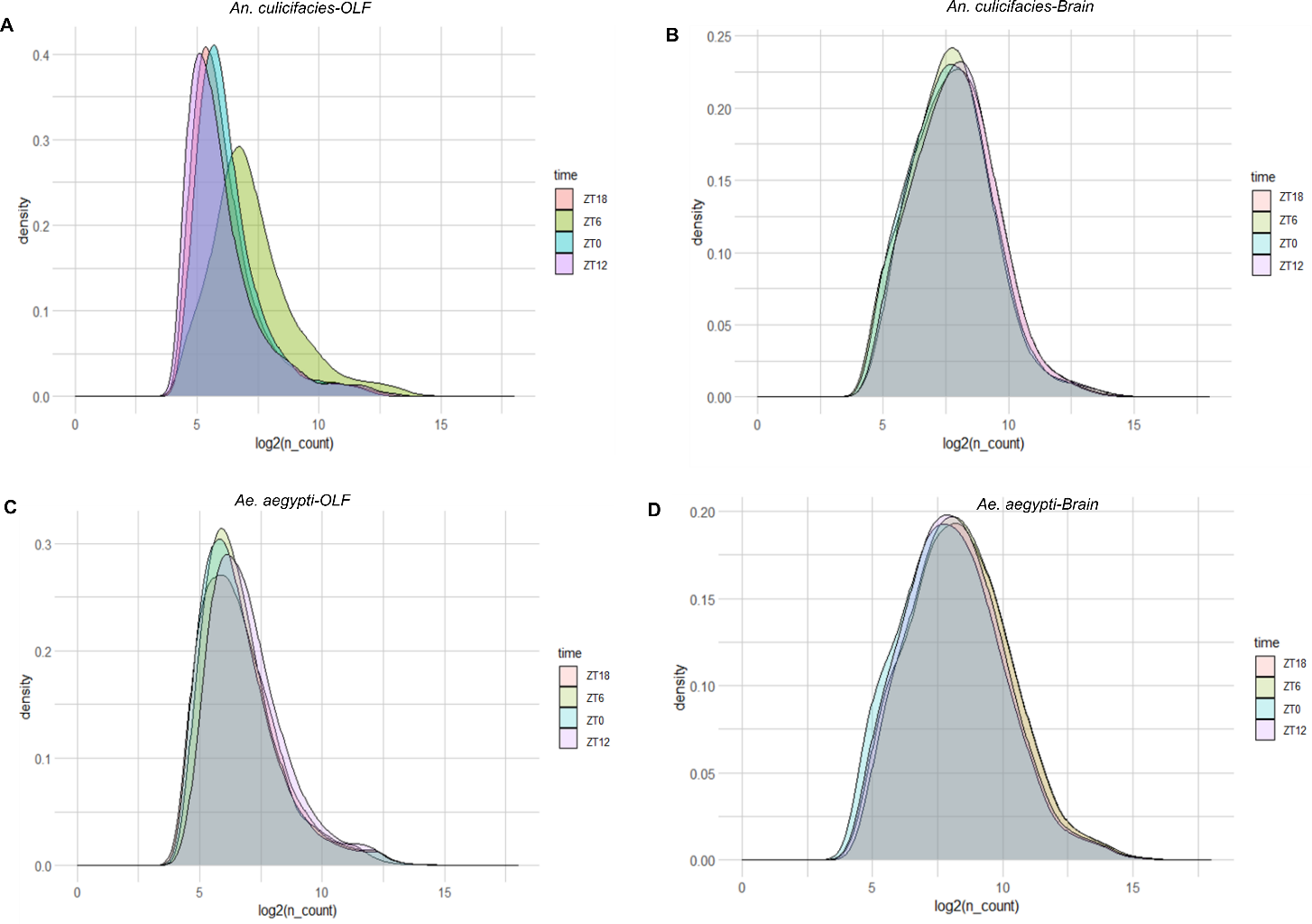

**Figure S1: Comparison of read density maps of peripheral olfactory and brain transcriptomes.** (A-B) Read density map of the circadian-dependent transcriptome data of peripheral olfactory (A) and brain (B) tissues of *An. culicifacies*. (C-D) Read density map of the circadian-dependent transcriptome data of peripheral olfactory (A) and brain (B) tissues of *Ae. aegypti*.

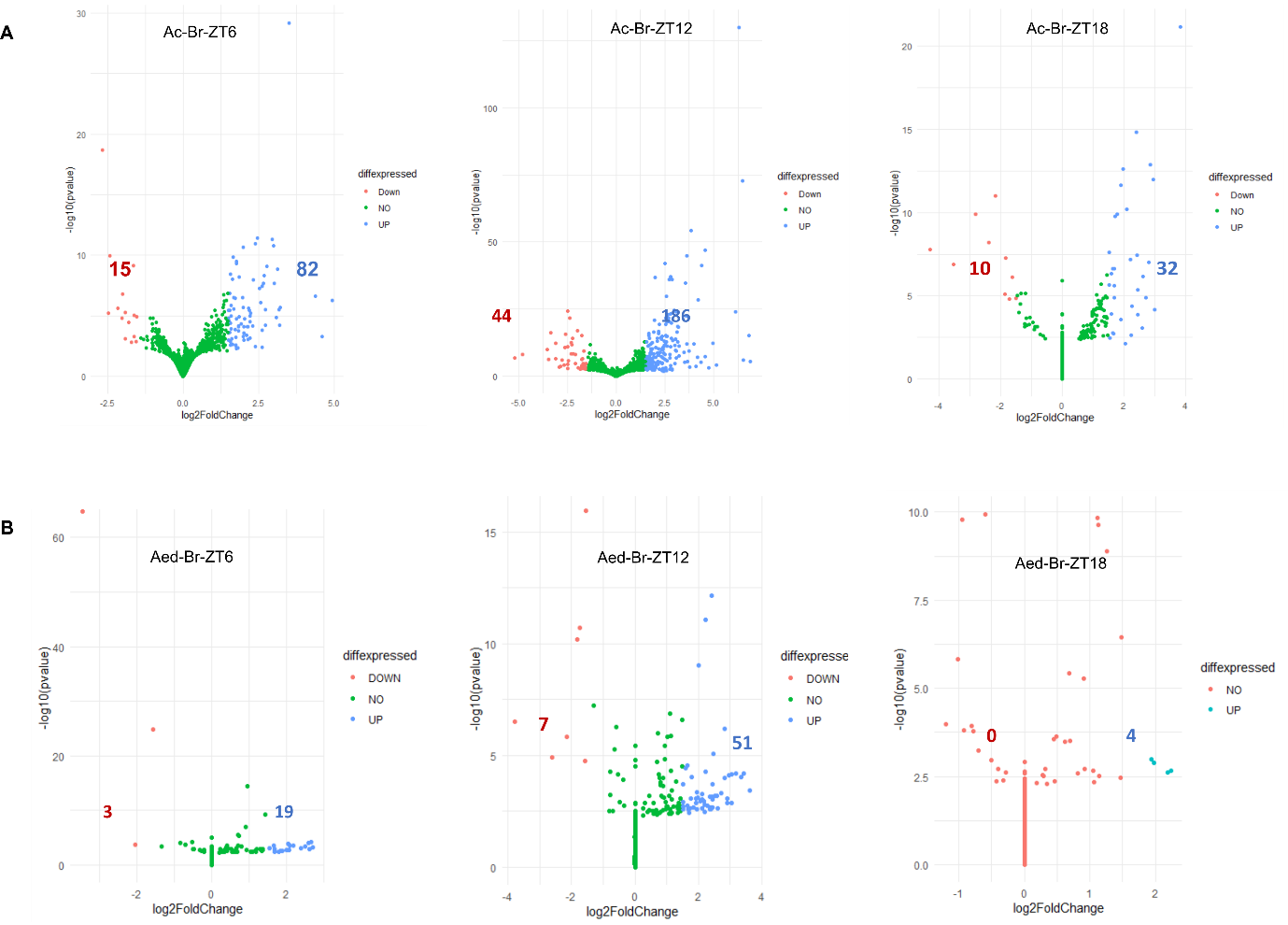

**Figure S2: Differential gene expression analysis of brain transcriptomes according to the circadian clock.** (A-B) Volcano plots of differentially expressed genes in the brain of *Anopheles culicifacies* (A) and *Aedes aegypti* (B). The number of upregulated and downregulated genes is mentioned in each respective plot.

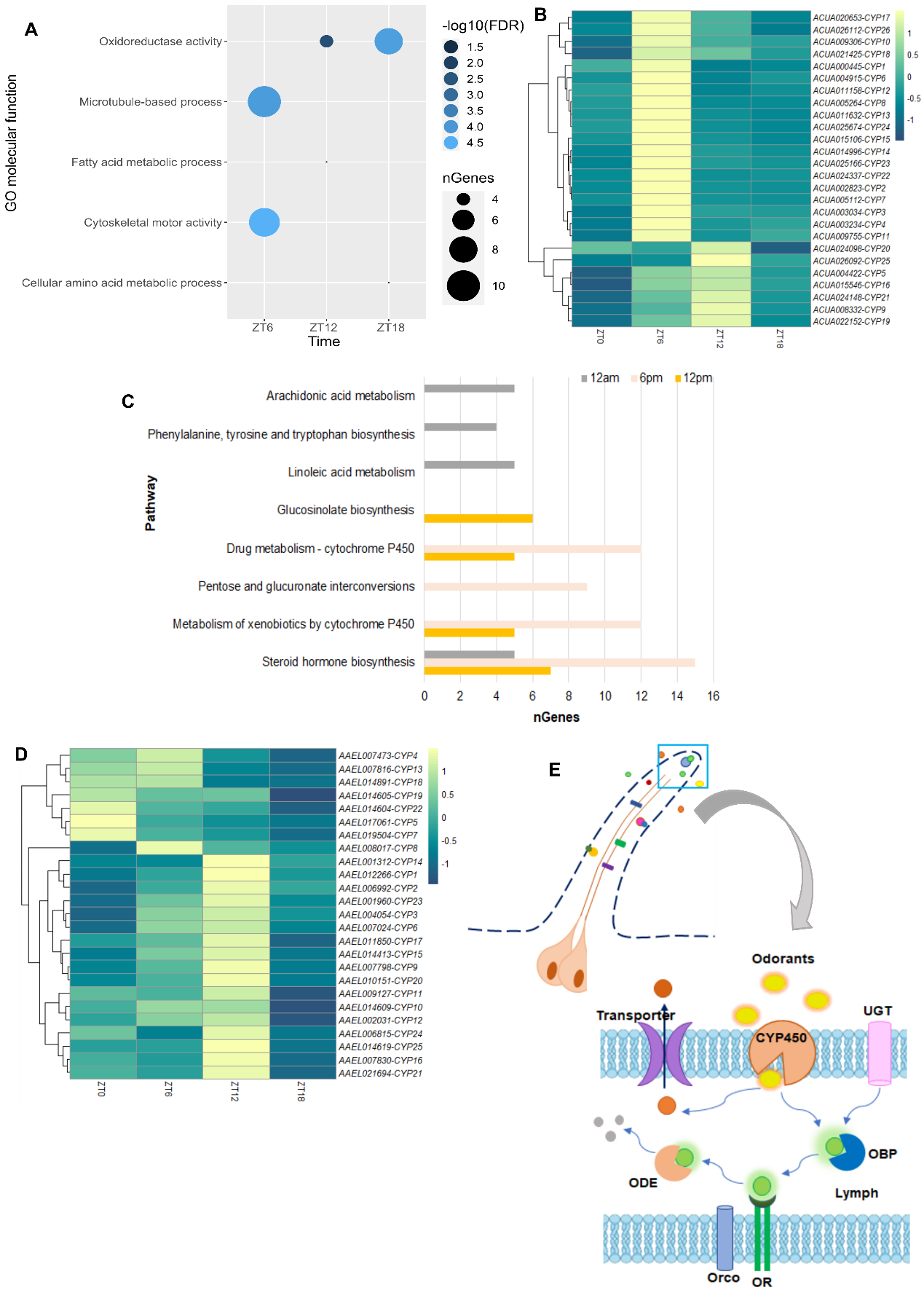

**Figure S3: Functional annotation analysis of differentially expressed olfactory genes of *Anopheles culicifacies*.** (A) Gene-set enrichment analysis of downregulated genes in the peripheral olfactory system. (B) Pathway enrichment analysis of differentially expressed olfactory genes of *An. culicifacies*. (C) Read-density dependent heatmap analysis of cytochrome P450 (CYP450) genes in *An. culicifacies*. (D) Read-density dependent heatmap analysis of cytochrome P450 (CYP450) genes in *Ae. aegypti. (*E) Hypothesized perireceptor events in the olfactory sensilla of mosquitoes. The upper panel indicates the structure of the sensilla. The right lower panel is the schematic representation of perireceptor events. Cumulative actions of two perireceptor proteins *i.e*., odorant binding proteins (OBPs) and xenobiotic metabolizing enzymes (XMEs), *i.e*., cytochrome P450 (CYP450) and UDP-glucosyltransferase (UGT) influence odorant detection by either mobilizing the odor molecules or modifying its chemical structure. Odorant degrading enzymes (ODE) degrade the odorant chemicals and also influence the resident time of the odorants within the sensillar lymph.

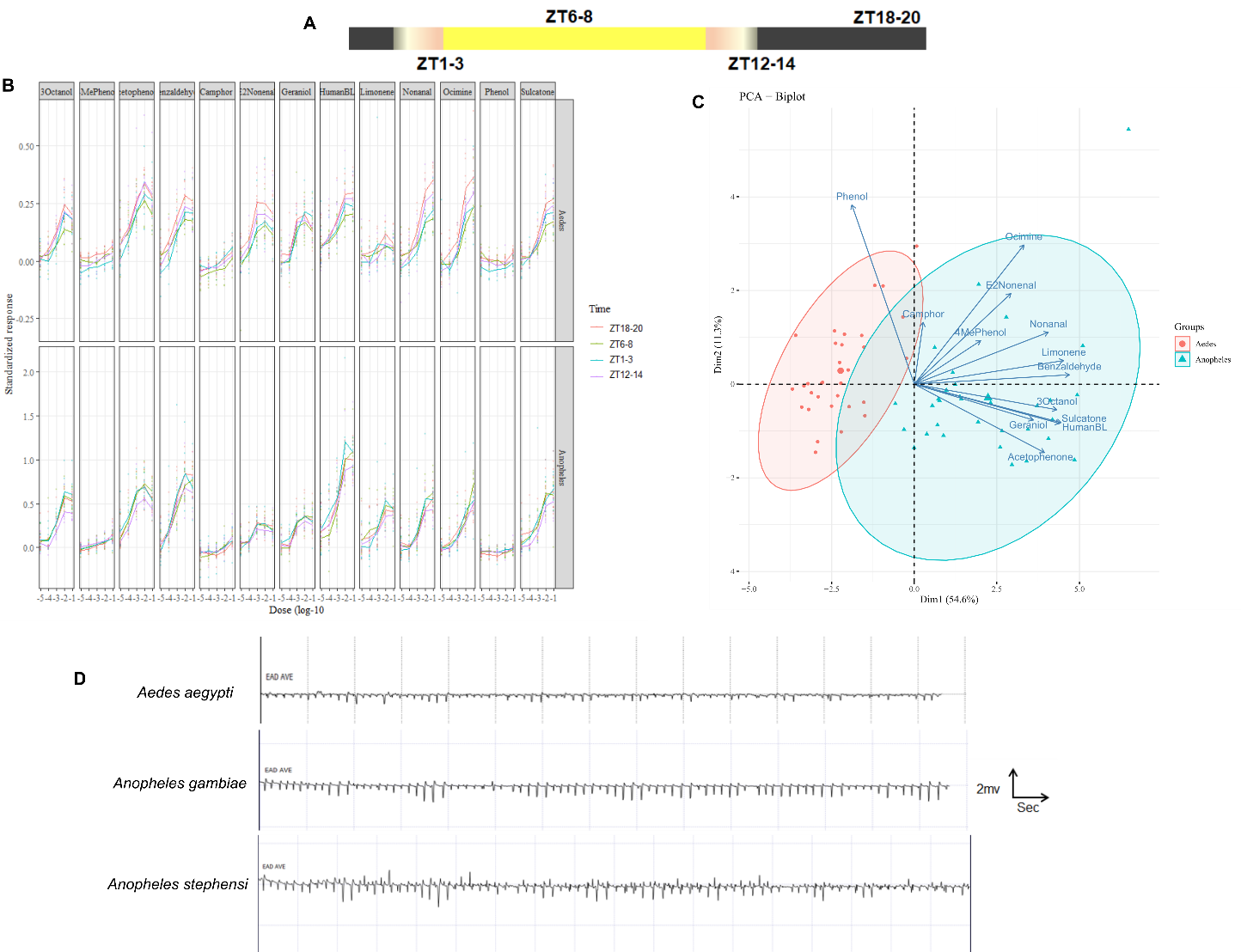

**Figure S4: Comparative electroantennographic response of diurnal and nocturnal mosquitoes.** (A) Schematic representation of the time points for EAG analysis. (B) Dose and time-of-day dependent relative EAG response in *An. gambiae* and *Ae. aegypti* against different volatile odorants. (C) Principal components analysis for making the connection between time of day, species, and compounds. (D) Comparative amplitude peaks (mV) of *Ae. aegypti*, *An. gambiae* and *An. stephensi* against increasing doses of 13 different odorant molecules, with solvent control (hexane) in between the two odorants. The order of the volatile odorants: camphor, 4-methylphenol, acetophenone, phenol, E2-nonenal, benzaldehyde, limonene, sulcatone, floral blend.

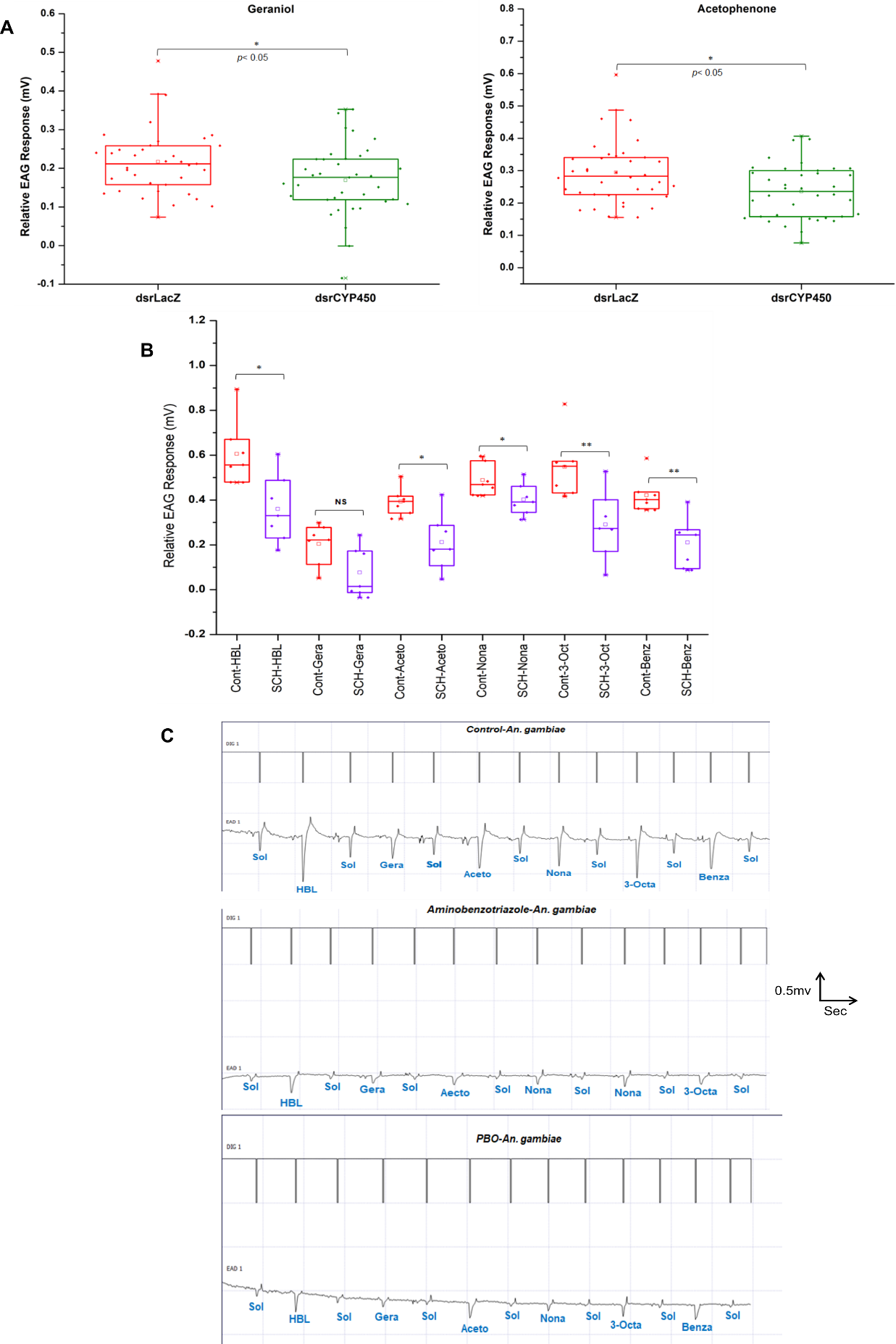

**Figure S5: Functional knock-down of CYP450 in mosquitoes.** (A) Relative EAG response of dsrLacZ and dsrCYP450 treated mosquitoes against two odorants, geraniol and acetophenone in *Ae. aegypti* (N=40). (B) EAG studies of naive vs inhibitor (Schinandrin A: CYP450 protein inhibitor) treated *An. gambiae*. (C) Representative EAG plots showing difference in the amplitude peaks (mV) in naive vs inhibitor-treated mosquitoes.

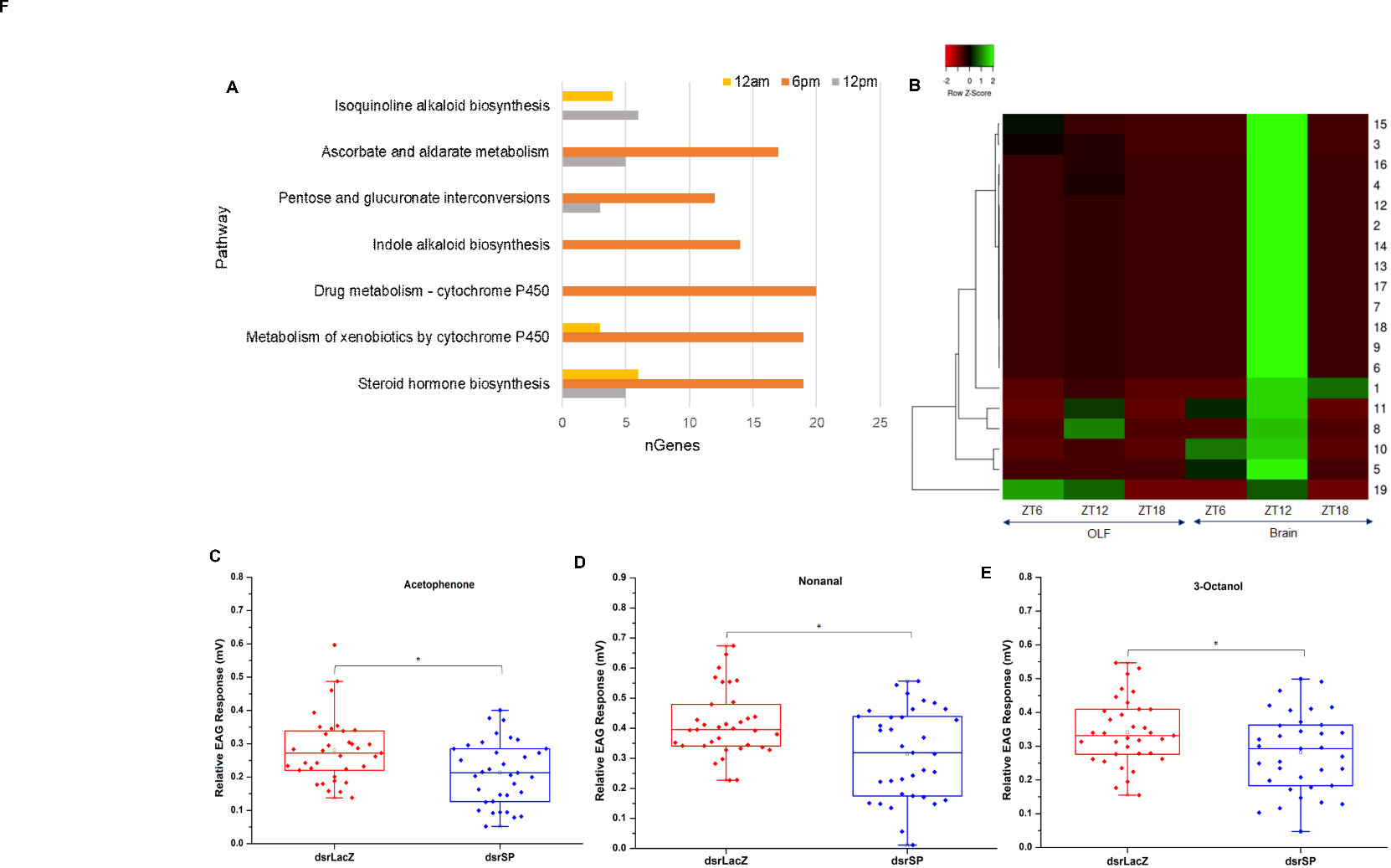

**Figure S6: Time-of-day dependent modulation of serine protease genes in the brain influence synaptic plasticity and olfactory sensitivity.** (A) Pathway enrichment analysis of upregulated brain transcripts. (B) Time-of-day dependent heatmap of serine-peptidase category of genes representing their significant enrichment in the brain of *Ae. aegypti* mosquitoes during ZT12. (C-E) Relative EAG response of naive (dsrLacZ injected) vs knock-down (dsrSP) *Ae. aegypti* to acetophenone, nonanal, and 3-octanol (N=40). Statistical significance was tested by Mann Whitney test for individual odorants and Significance values are indicated above bar graphs as asterisks (*), where p<=0.05.

**Table 1:** **Details of origin and the chemical nature of the volatile odorants tested**

| **Sl. No.** | **Compound Name** | **Possible Source** | **Chemical class** |
| --- | --- | --- | --- |
| 1. | Camphor | Floral | Terpenoid |
| 2. | Geraniol | Floral | Terpenoid |
| 3. | 4-Methyl Phenol | Human + Cattle | Phenolic |
| 4. | Acetophenone | Human + Cattle | Ketone |
| 5. | Phenol | Human + Cattle | Phenolic |
| 6. | Beta-Ocimine | Floral | Terpenoid |
| 7. | E2-Nonenal | Cattle | Aldehyde |
| 8. | Benzaldehyde | Human + Floral + Cattle | Aldehyde |
| 9. | Nonanl | Human + Floral + Cattle | Aldehyde |
| 10. | Limonene | Human + Floral + Cattle | Terpenoid |
| 11. | Sulcatone | Human + Floral + Cattle | Ketone |
| 12. | 3-Octanol | Human + Cattle | Alcohol |
| 13. | Human Blend |  |  |

Statistical analysis

To access if the temporal rhythms of mRNA expression showed an olfactory sensitivity to ecologically relevant volatile odorants a repeated measure ANOVA was carried out using dose and time as predictors in the model where individual identity of the mosquito (ID) was nested in dose for the two mosquito species. To conduct a comparative analysis between the two species a similar analysis was carried out as above while adding species identity as an additional predictor. If the time of day or its interaction terms were found to influence the antennal sensitivity a one-way ANOVA (within single species differences) or two-way ANOVA (between two species differences) followed by post hoc Tukey HSD test between the zeitgeber time points for each concentration of the volatile odorant (dose) was performed. To check the normality of the datasets, Shapiro Wilks test was carried out; or data were square root transformed to correct for non-normal distributions. All the above-mentioned analyses were conducted using the software R (version 4.3.1).

Supplementary data

Results

Individual species

Results of repeated measure ANOVA for EAG read outs of all compounds using dose, and time as predictors where individual identity (ID) is nested in dose for ***Anopheles****.* All the compounds where ‘Time’ or its interaction terms were found to be significant are highlighted in yellow.

```
Camphor

Error: ID
          Df Sum Sq  Mean Sq F value Pr(>F)
Time       3 0.0233 0.007755   0.301  0.824
Residuals 28 0.7214 0.025764

Error: ID:Dose
           Df Sum Sq Mean Sq F value Pr(>F)
Dose        4 0.6882 0.17205  57.296 <2e-16 ***
Dose:Time  12 0.0537 0.00448   1.491  0.138
Residuals 112 0.3363 0.00300
---
Signif. codes:  0 ‘***’ 0.001 ‘**’ 0.01 ‘*’ 0.05 ‘.’ 0.1 ‘ ’ 1

Geraniol

Error: ID
          Df Sum Sq Mean Sq F value Pr(>F)
Time       3 0.0631 0.02102   0.467  0.708
Residuals 28 1.2603 0.04501

Error: ID:Dose
           Df Sum Sq Mean Sq F value Pr(>F)
Dose        4 2.9673  0.7418 133.778 <2e-16 ***
Dose:Time  12 0.0845  0.0070   1.271  0.246
Residuals 112 0.6211  0.0055
---
Signif. codes:  0 ‘***’ 0.001 ‘**’ 0.01 ‘*’ 0.05 ‘.’ 0.1 ‘ ’ 1

4MePhenol

Error: ID
          Df Sum Sq  Mean Sq F value Pr(>F)
Time       3 0.0164 0.005481   0.482  0.698
Residuals 28 0.3186 0.011378

Error: ID:Dose
           Df Sum Sq Mean Sq F value Pr(>F)
Dose        4 0.3822 0.09554  56.341 <2e-16 ***
Dose:Time  12 0.0289 0.00241   1.419  0.168
Residuals 112 0.1899 0.00170
---
Signif. codes:  0 ‘***’ 0.001 ‘**’ 0.01 ‘*’ 0.05 ‘.’ 0.1 ‘ ’ 1

Acetophenone

Error: ID
          Df Sum Sq Mean Sq F value Pr(>F)
Time       3 0.4593 0.15310   2.282  0.101
Residuals 28 1.8782 0.06708

Error: ID:Dose
           Df Sum Sq Mean Sq F value Pr(>F)
Dose        4  6.432  1.6081  90.727 <2e-16 ***
Dose:Time  12  0.126  0.0105   0.595  0.843
Residuals 112  1.985  0.0177
---
Signif. codes:  0 ‘***’ 0.001 ‘**’ 0.01 ‘*’ 0.05 ‘.’ 0.1 ‘ ’ 1

Phenol

Error: ID
          Df  Sum Sq  Mean Sq F value Pr(>F)
Time       3 0.01819 0.006065   0.625  0.605
Residuals 28 0.27162 0.009701

Error: ID:Dose
           Df  Sum Sq  Mean Sq F value   Pr(>F)
Dose        4 0.05661 0.014153  23.587 3.66e-14 ***
Dose:Time  12 0.02009 0.001675   2.791  0.00229 **
Residuals 112 0.06720 0.000600
---
Signif. codes:  0 ‘***’ 0.001 ‘**’ 0.01 ‘*’ 0.05 ‘.’ 0.1 ‘ ’ 1

Ocimine

Error: ID
          Df Sum Sq Mean Sq F value Pr(>F)
Time       3 0.0954 0.03179   0.674  0.575
Residuals 28 1.3200 0.04714

Error: ID:Dose
           Df Sum Sq Mean Sq F value Pr(>F)
Dose        4  7.251  1.8127  99.374 <2e-16 ***
Dose:Time  12  0.476  0.0396   2.174 0.0176 *
Residuals 112  2.043  0.0182
---
Signif. codes:  0 ‘***’ 0.001 ‘**’ 0.01 ‘*’ 0.05 ‘.’ 0.1 ‘ ’ 1


E2Nonenal

Error: ID
          Df Sum Sq Mean Sq F value Pr(>F)
Time       3 0.0365 0.01218   0.423  0.738
Residuals 28 0.8066 0.02881

Error: ID:Dose
           Df Sum Sq Mean Sq F value Pr(>F)
Dose        4 1.3338  0.3335  59.976 <2e-16 ***
Dose:Time  12 0.0582  0.0048   0.872  0.577
Residuals 112 0.6227  0.0056
---
Signif. codes:  0 ‘***’ 0.001 ‘**’ 0.01 ‘*’ 0.05 ‘.’ 0.1 ‘ ’ 1


Benzaldehyde

Error: ID
          Df Sum Sq Mean Sq F value Pr(>F)
Time       3  0.225 0.07511   0.593  0.625
Residuals 28  3.549 0.12675

Error: ID:Dose
           Df Sum Sq Mean Sq F value Pr(>F)
Dose        4 13.747   3.437 110.144 <2e-16 ***
Dose:Time  12  0.345   0.029   0.921  0.529
Residuals 112  3.495   0.031
---
Signif. codes:  0 ‘***’ 0.001 ‘**’ 0.01 ‘*’ 0.05 ‘.’ 0.1 ‘ ’ 1

Nonanal

Error: ID
          Df Sum Sq Mean Sq F value Pr(>F)
Time       3 0.1104 0.03682   0.492  0.691
Residuals 28 2.0946 0.07481

Error: ID:Dose
           Df Sum Sq Mean Sq F value Pr(>F)
Dose        4  8.576  2.1440 108.486 <2e-16 ***
Dose:Time  12  0.224  0.0187   0.944  0.506
Residuals 112  2.213  0.0198
---
Signif. codes:  0 ‘***’ 0.001 ‘**’ 0.01 ‘*’ 0.05 ‘.’ 0.1 ‘ ’ 1

Limonene

Error: ID
          Df Sum Sq Mean Sq F value Pr(>F)
Time       3 0.1355 0.04517   0.475  0.702
Residuals 28 2.6621 0.09508

Error: ID:Dose
           Df Sum Sq Mean Sq F value Pr(>F)
Dose        4  4.173  1.0432  37.643 <2e-16 ***
Dose:Time  12  0.248  0.0206   0.745  0.705
Residuals 112  3.104  0.0277
---
Signif. codes:  0 ‘***’ 0.001 ‘**’ 0.01 ‘*’ 0.05 ‘.’ 0.1 ‘ ’ 1

Sulcatone

Error: ID
          Df Sum Sq Mean Sq F value Pr(>F)
Time       3 0.3253 0.10843   1.413   0.26
Residuals 28 2.1481 0.07672

Error: ID:Dose
           Df Sum Sq Mean Sq F value Pr(>F)
Dose        4  8.131  2.0328  67.504 <2e-16 ***
Dose:Time  12  0.145  0.0121   0.401  0.961
Residuals 112  3.373  0.0301
---
Signif. codes:  0 ‘***’ 0.001 ‘**’ 0.01 ‘*’ 0.05 ‘.’ 0.1 ‘ ’ 1

3Octanol

Error: ID
          Df Sum Sq Mean Sq F value Pr(>F)
Time       3 0.3933 0.13109   3.067 0.0441 *
Residuals 28 1.1966 0.04274
---
Signif. codes:  0 ‘***’ 0.001 ‘**’ 0.01 ‘*’ 0.05 ‘.’ 0.1 ‘ ’ 1

Error: ID:Dose
           Df Sum Sq Mean Sq F value Pr(>F)
Dose        4  6.909  1.7273 158.423 <2e-16 ***
Dose:Time  12  0.126  0.0105   0.966  0.485
Residuals 112  1.221  0.0109
---
Signif. codes:  0 ‘***’ 0.001 ‘**’ 0.01 ‘*’ 0.05 ‘.’ 0.1 ‘ ’ 1

Human Blend


Error: ID
          Df Sum Sq Mean Sq F value Pr(>F)
Time       3  0.502  0.1674   0.937  0.436
Residuals 28  5.003  0.1787

Error: ID:Dose
           Df Sum Sq Mean Sq F value Pr(>F)
Dose        4 21.171   5.293 149.632 <2e-16 ***
Dose:Time  12  0.457   0.038   1.077  0.386
Residuals 112  3.962   0.035
---
Signif. codes:  0 ‘***’ 0.001 ‘**’ 0.01 ‘*’ 0.05 ‘.’ 0.1 ‘ ’ 1
```

Results of one-way ANOVA and post hoc Tukey HSD test for the EAG readouts for Phenol, β Ocimene and 3 Octanol to figure out the difference between different time points at each concentration/dose. Compounds showing differences between the time points were highlighted by blue, concentration by green and time points pair by grey.

```
#Phenol

Model used: Phenol ~ Time

#10^5 concentration

            Df  Sum Sq  Mean Sq F value Pr(>F)
Time         3 0.00587 0.001955   0.811  0.498
Residuals   28 0.06746 0.002409

#10^4 concentration

            Df  Sum Sq  Mean Sq F value Pr(>F)
Time         3 0.00948 0.003160   1.194   0.33
Residuals   28 0.07413 0.002648

#10^3 concentration

            Df  Sum Sq  Mean Sq F value Pr(>F)
Time         3 0.01044 0.003482   1.191  0.331
Residuals   28 0.08186 0.002924


#10^2 concentration

            Df  Sum Sq  Mean Sq F value Pr(>F)
Time         3 0.00924 0.003080   1.679  0.194
Residuals   28 0.05137 0.001835


#10^1 concentration

            Df  Sum Sq  Mean Sq F value Pr(>F)
Time         3 0.00326 0.001086   0.475  0.702
Residuals   28 0.06400 0.002286


#Ocimene

#10^5 concentration

            Df  Sum Sq  Mean Sq F value Pr(>F)
Time         3 0.00485 0.001616   0.382  0.767
Residuals   28 0.11843 0.004230

#10^4 concentration

            Df  Sum Sq  Mean Sq F value Pr(>F)
Time         3 0.02525 0.008417    0.78  0.515
Residuals   28 0.30227 0.010795

#10^3 concentration

            Df Sum Sq Mean Sq F value Pr(>F)
Time         3 0.0419 0.01397   0.414  0.744
Residuals   28 0.9451 0.03375

#10^2 concentration

            Df Sum Sq Mean Sq F value Pr(>F)
Time         3 0.0837 0.02792   0.769  0.521
Residuals   28 1.0161 0.03629

#10^1 concentration
            Df Sum Sq Mean Sq F value Pr(>F)
Time         3 0.4154 0.13847   3.952 0.0181 *
Residuals   28 0.9811 0.03504
---
Signif. codes:  0 ‘***’ 0.001 ‘**’ 0.01 ‘*’ 0.05 ‘.’ 0.1 ‘ ’ 1

Tukey multiple comparisons of means
    95% family-wise confidence level

$Time
                     diff         lwr        upr     p adj
2.30pm-2.30am  0.31034206  0.05480373 0.56588039 0.0127655 *
8.30am-2.30am  0.11159206 -0.14394627 0.36713039 0.6366311
8.30pm-2.30am  0.08287006 -0.17266827 0.33840839 0.8123773
8.30am-2.30pm -0.19875000 -0.45428833 0.05678833 0.1703927
8.30pm-2.30pm -0.22747200 -0.48301033 0.02806633 0.0943306
8.30pm-8.30am -0.02872200 -0.28426033 0.22681633 0.9897506


#3Octanol

#10^5 concentration

           Df Sum Sq  Mean Sq F value Pr(>F)
Time         3 0.0060 0.001996   0.146  0.931
Residuals   28 0.3823 0.013652

#10^4 concentration

            Df Sum Sq Mean Sq F value Pr(>F)
Time         3 0.0398 0.01326   0.727  0.544
Residuals   28 0.5107 0.01824

#10^3 concentration

           Df Sum Sq Mean Sq F value Pr(>F)
Time         3 0.0612 0.02039   1.222   0.32
Residuals   28 0.4672 0.01669

#10^2 concentration

         Df Sum Sq Mean Sq F value Pr(>F)
Time         3 0.2243 0.07477   3.737 0.0224 *
Residuals   28 0.5603 0.02001
---
Signif. codes:  0 ‘***’ 0.001 ‘**’ 0.01 ‘*’ 0.05 ‘.’ 0.1 ‘ ’ 1

Tukey multiple comparisons of means
    95% family-wise confidence level

$Time
                     diff        lwr         upr     p adj
2.30pm-2.30am  0.01963294 -0.1734741  0.21273994 0.9923610
8.30am-2.30am  0.07088294 -0.1222241  0.26398994 0.7493755
8.30pm-2.30am -0.15371431 -0.3468213  0.03939269 0.1554983
8.30am-2.30pm  0.05125000 -0.1418570  0.24435700 0.8864274
8.30pm-2.30pm -0.17334725 -0.3664543  0.01975975 0.0904769
8.30pm-8.30am -0.22459725 -0.4177043 -0.03149025 0.0179312*


#10^1 concentration


           Df Sum Sq Mean Sq F value Pr(>F)
Time         3 0.1884 0.06279   3.536 0.0274 *
Residuals   28 0.4973 0.01776
---
Signif. codes:  0 ‘***’ 0.001 ‘**’ 0.01 ‘*’ 0.05 ‘.’ 0.1 ‘ ’ 1

Tukey multiple comparisons of means
    95% family-wise confidence level

$Time
                     diff        lwr         upr     p adj
2.30pm-2.30am  0.01944456 -0.1624803  0.20136940 0.9911540
8.30am-2.30am  0.07194456 -0.1099803  0.25386940 0.7044824
8.30pm-2.30am -0.13598044 -0.3179053  0.04594440 0.1975632
8.30am-2.30pm  0.05250000 -0.1294248  0.23442484 0.8593034
8.30pm-2.30pm -0.15542500 -0.3373498  0.02649984 0.1146894
8.30pm-8.30am -0.20792500 -0.3898498 -0.02600016 0.0204470 *
```

Results of repeated measure ANOVA for EAG readouts of all compounds using dose, and time as predictors where individual identity (ID) is nested in dose for ***Aedes***. All the compounds where ‘Time’ or its interaction terms were found to be significant are highlighted in yellow.

```
Camphor

Error: ID
          Df  Sum Sq  Mean Sq F value Pr(>F)
Time       3 0.03205 0.010684   2.298 0.0992 .
Residuals 28 0.13016 0.004649
---
Signif. codes:  0 ‘***’ 0.001 ‘**’ 0.01 ‘*’ 0.05 ‘.’ 0.1 ‘ ’ 1

Error: ID:Dose
           Df  Sum Sq Mean Sq F value Pr(>F)
Dose        4 0.13118 0.03280  47.330 <2e-16 ***
Dose:Time  12 0.00688 0.00057   0.828  0.622
Residuals 112 0.07761 0.00069
---
Signif. codes:  0 ‘***’ 0.001 ‘**’ 0.01 ‘*’ 0.05 ‘.’ 0.1 ‘ ’ 1

Geraniol

Error: ID
          Df  Sum Sq  Mean Sq F value Pr(>F)
Time       3 0.01623 0.005410   0.658  0.584
Residuals 28 0.23004 0.008216

Error: ID:Dose
           Df Sum Sq Mean Sq F value Pr(>F)
Dose        4 1.0067 0.25167  77.265 <2e-16 ***
Dose:Time  12 0.0477 0.00398   1.221  0.278
Residuals 112 0.3648 0.00326
---
Signif. codes:  0 ‘***’ 0.001 ‘**’ 0.01 ‘*’ 0.05 ‘.’ 0.1 ‘ ’ 1

4MePhenol
```

```
Error: ID
          Df  Sum Sq  Mean Sq F value   Pr(>F)
Time       3 0.06153 0.020508   8.067 0.000499 ***
Residuals 28 0.07118 0.002542
---
Signif. codes:  0 ‘***’ 0.001 ‘**’ 0.01 ‘*’ 0.05 ‘.’ 0.1 ‘ ’ 1

Error: ID:Dose
           Df  Sum Sq  Mean Sq F value Pr(>F)
Dose        4 0.05153 0.012884  40.192 <2e-16 ***
Dose:Time  12 0.00691 0.000576   1.797 0.0568 .
Residuals 112 0.03590 0.000321
---
Signif. codes:  0 ‘***’ 0.001 ‘**’ 0.01 ‘*’ 0.05 ‘.’ 0.1 ‘ ’ 1

Acetophenone

Error: ID
          Df Sum Sq Mean Sq F value Pr(>F)
Time       3 0.0792 0.02641   0.926  0.441
Residuals 28 0.7982 0.02851

Error: ID:Dose
           Df Sum Sq Mean Sq F value Pr(>F)
Dose        4 1.2898  0.3225 129.799 <2e-16 ***
Dose:Time  12 0.0471  0.0039   1.581  0.107
Residuals 112 0.2782  0.0025
---
Signif. codes:  0 ‘***’ 0.001 ‘**’ 0.01 ‘*’ 0.05 ‘.’ 0.1 ‘ ’ 1


Phenol

Error: ID
          Df  Sum Sq  Mean Sq F value Pr(>F)
Time       3 0.03987 0.013290   4.373  0.012 *
Residuals 28 0.08510 0.003039
---
Signif. codes:  0 ‘***’ 0.001 ‘**’ 0.01 ‘*’ 0.05 ‘.’ 0.1 ‘ ’ 1

Error: ID:Dose
           Df  Sum Sq  Mean Sq F value   Pr(>F)
Dose        4 0.02017 0.005043   7.569 1.96e-05 ***
Dose:Time  12 0.01337 0.001114   1.673   0.0822 .
Residuals 112 0.07462 0.000666
---
Signif. codes:  0 ‘***’ 0.001 ‘**’ 0.01 ‘*’ 0.05 ‘.’ 0.1 ‘ ’ 1

Ocimine

Error: ID
          Df Sum Sq Mean Sq F value Pr(>F)
Time       3 0.1582 0.05274   4.009 0.0171 *
Residuals 28 0.3683 0.01315
---
Signif. codes:  0 ‘***’ 0.001 ‘**’ 0.01 ‘*’ 0.05 ‘.’ 0.1 ‘ ’ 1

Error: ID:Dose
           Df Sum Sq Mean Sq F value Pr(>F)
Dose        4 2.4615  0.6154  97.541 <2e-16 ***
Dose:Time  12 0.0897  0.0075   1.186  0.302
Residuals 112 0.7066  0.0063
---
Signif. codes:  0 ‘***’ 0.001 ‘**’ 0.01 ‘*’ 0.05 ‘.’ 0.1 ‘ ’ 1

E2Nonenal
```

```
Error: ID
          Df Sum Sq Mean Sq F value Pr(>F)
Time       3 0.1885 0.06285   3.032 0.0458 *
Residuals 28 0.5805 0.02073
---
Signif. codes:  0 ‘***’ 0.001 ‘**’ 0.01 ‘*’ 0.05 ‘.’ 0.1 ‘ ’ 1

Error: ID:Dose
           Df Sum Sq Mean Sq F value Pr(>F)
Dose        4 0.8315 0.20789  53.680 <2e-16 ***
Dose:Time  12 0.0188 0.00157   0.405  0.959
Residuals 112 0.4337 0.00387
---
Signif. codes:  0 ‘***’ 0.001 ‘**’ 0.01 ‘*’ 0.05 ‘.’ 0.1 ‘ ’ 1

Benzaldehyde

Error: ID
          Df Sum Sq Mean Sq F value Pr(>F)
Time       3 0.1144 0.03814    3.82 0.0206 *
Residuals 28 0.2796 0.00998
---
Signif. codes:  0 ‘***’ 0.001 ‘**’ 0.01 ‘*’ 0.05 ‘.’ 0.1 ‘ ’ 1

Error: ID:Dose
           Df Sum Sq Mean Sq F value Pr(>F)
Dose        4 1.3431  0.3358 143.906 <2e-16 ***
Dose:Time  12 0.0537  0.0045   1.917 0.0395 *
Residuals 112 0.2613  0.0023
---
Signif. codes:  0 ‘***’ 0.001 ‘**’ 0.01 ‘*’ 0.05 ‘.’ 0.1 ‘ ’ 1

Nonanal

Error: ID
          Df Sum Sq Mean Sq F value  Pr(>F)
Time       3 0.1863 0.06210   4.948 0.00701 **
Residuals 28 0.3514 0.01255
---
Signif. codes:  0 ‘***’ 0.001 ‘**’ 0.01 ‘*’ 0.05 ‘.’ 0.1 ‘ ’ 1

Error: ID:Dose
           Df Sum Sq Mean Sq F value  Pr(>F)
Dose        4 1.8143  0.4536 238.467 < 2e-16 ***
Dose:Time  12 0.1165  0.0097   5.106 9.5e-07 ***
Residuals 112 0.2130  0.0019
---
Signif. codes:  0 ‘***’ 0.001 ‘**’ 0.01 ‘*’ 0.05 ‘.’ 0.1 ‘ ’ 1

Limonene

Error: ID
          Df  Sum Sq  Mean Sq F value Pr(>F)
Time       3 0.01997 0.006658   0.609  0.615
Residuals 28 0.30594 0.010926

Error: ID:Dose
           Df Sum Sq  Mean Sq F value   Pr(>F)
Dose        4 0.1065 0.026637   7.479 2.24e-05 ***
Dose:Time  12 0.0339 0.002823   0.793    0.657
Residuals 112 0.3989 0.003561
---
Signif. codes:  0 ‘***’ 0.001 ‘**’ 0.01 ‘*’ 0.05 ‘.’ 0.1 ‘ ’ 1

Sulcatone

Error: ID
          Df  Sum Sq  Mean Sq F value Pr(>F)
Time       3 0.05155 0.017184   1.965  0.142
Residuals 28 0.24483 0.008744

Error: ID:Dose
           Df Sum Sq Mean Sq F value Pr(>F)
Dose        4  1.359  0.3397 178.559 <2e-16 ***
Dose:Time  12  0.046  0.0038   2.014 0.0291 *
Residuals 112  0.213  0.0019
---
Signif. codes:  0 ‘***’ 0.001 ‘**’ 0.01 ‘*’ 0.05 ‘.’ 0.1 ‘ ’ 1

3Octanol

Error: ID
          Df  Sum Sq  Mean Sq F value Pr(>F)
Time       3 0.05938 0.019794   2.827 0.0567 .
Residuals 28 0.19607 0.007003
---
Signif. codes:  0 ‘***’ 0.001 ‘**’ 0.01 ‘*’ 0.05 ‘.’ 0.1 ‘ ’ 1

Error: ID:Dose
           Df Sum Sq Mean Sq F value  Pr(>F)
Dose        4 0.9096 0.22741 138.843 < 2e-16 ***
Dose:Time  12 0.0523 0.00436   2.661 0.00354 **
Residuals 112 0.1834 0.00164
---
Signif. codes:  0 ‘***’ 0.001 ‘**’ 0.01 ‘*’ 0.05 ‘.’ 0.1 ‘ ’ 1


Human Blend

Error: ID
          Df Sum Sq Mean Sq F value Pr(>F)
Time       3 0.0623 0.02078   0.917  0.446
Residuals 28 0.6349 0.02268

Error: ID:Dose
           Df Sum Sq Mean Sq F value Pr(>F)
Dose        4 0.9731 0.24328 126.836 <2e-16 ***
Dose:Time  12 0.0359 0.00300   1.562  0.113
Residuals 112 0.2148 0.00192
---
Signif. codes:  0 ‘***’ 0.001 ‘**’ 0.01 ‘*’ 0.05 ‘.’ 0.1 ‘ ’ 1
```

Results of one-way ANOVA and post hoc Tukey HSD test for the EAG readouts for 4MePhenol, Ocimene, E2Nonenal, Nonanal, Benzaldehyde, Sulcatone and 3Octanol to figure out the difference between different time points at each concentration/dose. Compounds showing differences between the time points were highlighted by blue, concentration by green and time points pair by grey.

```
#4MePhenol

#10^5 concentration

            Df  Sum Sq  Mean Sq F value  Pr(>F)
Time         3 0.02032 0.006772   6.791 0.00139 **
Residuals   28 0.02792 0.000997
---
Signif. codes:  0 ‘***’ 0.001 ‘**’ 0.01 ‘*’ 0.05 ‘.’ 0.1 ‘ ’ 1

Tukey multiple comparisons of means
    95% family-wise confidence level

$Time
                     diff         lwr          upr     p adj
2.30pm-2.30am -0.01184638 -0.05495665  0.031263899 0.8757736
8.30am-2.30am -0.06566545 -0.10877572 -0.022555176 0.0014796 **
8.30pm-2.30am -0.03679081 -0.07990109  0.006319461 0.1152597
8.30am-2.30pm -0.05381907 -0.09692935 -0.010708801 0.0101624 *
8.30pm-2.30pm -0.02494444 -0.06805471  0.018165836 0.4059526
8.30pm-8.30am  0.02887464 -0.01423564  0.071984911 0.2815332

#10^4 concentration

            Df   Sum Sq   Mean Sq F value Pr(>F)
Time         3 0.009194 0.0030646   4.531 0.0104 *
Residuals   28 0.018939 0.0006764
---
Signif. codes:  0 ‘***’ 0.001 ‘**’ 0.01 ‘*’ 0.05 ‘.’ 0.1 ‘ ’ 1

Tukey multiple comparisons of means
    95% family-wise confidence level

$Time
                      diff         lwr           upr     p adj
2.30pm-2.30am -0.018700913 -0.05420529  1.680347e-02 0.4870917
8.30am-2.30am -0.044371500 -0.07987588 -8.867118e-03 0.0100706 *
8.30pm-2.30am -0.035479438 -0.07098382  2.494493e-05 0.0502114
8.30am-2.30pm -0.025670587 -0.06117497  9.833795e-03 0.2217188
8.30pm-2.30pm -0.016778525 -0.05228291  1.872586e-02 0.5765319
8.30pm-8.30am  0.008892062 -0.02661232  4.439644e-02 0.9023874

#10^3 concentration

            Df  Sum Sq  Mean Sq F value  Pr(>F)
Time         3 0.01302 0.004339   6.889 0.00128 **
Residuals   28 0.01763 0.000630
---
Signif. codes:  0 ‘***’ 0.001 ‘**’ 0.01 ‘*’ 0.05 ‘.’ 0.1 ‘ ’ 1

Tukey multiple comparisons of means
    95% family-wise confidence level

$Time
                      diff          lwr          upr     p adj
2.30pm-2.30am -0.037476500 -0.071736083 -0.003216917 0.0279985 *
8.30am-2.30am -0.055882750 -0.090142333 -0.021623167 0.0006758 ***
8.30pm-2.30am -0.028565313 -0.062824896  0.005694271 0.1279254
8.30am-2.30pm -0.018406250 -0.052665833  0.015853333 0.4701537
8.30pm-2.30pm  0.008911187 -0.025348396  0.043170771 0.8922155
8.30pm-8.30am  0.027317438 -0.006942146  0.061577021 0.1544296

#10^2 concentration

            Df   Sum Sq   Mean Sq F value Pr(>F)
Time         3 0.008989 0.0029963   4.418 0.0115 *
Residuals   28 0.018990 0.0006782
---
Signif. codes:  0 ‘***’ 0.001 ‘**’ 0.01 ‘*’ 0.05 ‘.’ 0.1 ‘ ’ 1

Tukey multiple comparisons of means
    95% family-wise confidence level

$Time
                      diff          lwr          upr     p adj
2.30pm-2.30am -0.004574375 -0.040126974  0.030978224 0.9848085
8.30am-2.30am -0.043142250 -0.078694849 -0.007589651 0.0128499*
8.30pm-2.30am -0.015184313 -0.050736911  0.020368286 0.6526150
8.30am-2.30pm -0.038567875 -0.074120474 -0.003015276 0.0296537*
8.30pm-2.30pm -0.010609937 -0.046162536  0.024942661 0.8469555
8.30pm-8.30am  0.027957937 -0.007594661  0.063510536 0.1632255

#10^1 concentration

            Df  Sum Sq  Mean Sq F value  Pr(>F)
Time         3 0.01692 0.005642   6.695 0.00151 **
Residuals   28 0.02360 0.000843
---
Signif. codes:  0 ‘***’ 0.001 ‘**’ 0.01 ‘*’ 0.05 ‘.’ 0.1 ‘ ’ 1

Tukey multiple comparisons of means
    95% family-wise confidence level

$Time
                      diff         lwr          upr     p adj
2.30pm-2.30am -0.034227625 -0.07385686  0.005401612 0.1090467
8.30am-2.30am -0.064964875 -0.10459411 -0.025335638 0.0006367***
8.30pm-2.30am -0.030949438 -0.07057867  0.008679800 0.1676975
8.30am-2.30pm -0.030737250 -0.07036649  0.008891987 0.1722188
8.30pm-2.30pm  0.003278188 -0.03635105  0.042907425 0.9958410
8.30pm-8.30am  0.034015438 -0.00561380  0.073644675 0.1122456


#Ocimene

#10^5 concentration

           Df  Sum Sq  Mean Sq F value Pr(>F)
Time         3 0.00694 0.002313   0.474  0.703
Residuals   28 0.13664 0.004880

#10^4 concentration

            Df  Sum Sq  Mean Sq F value Pr(>F)
Time         3 0.02662 0.008874   3.821 0.0206 *
Residuals   28 0.06503 0.002323
---
Signif. codes:  0 ‘***’ 0.001 ‘**’ 0.01 ‘*’ 0.05 ‘.’ 0.1 ‘ ’ 1

Tukey multiple comparisons of means
    95% family-wise confidence level

$Time
                      diff         lwr          upr     p adj
2.30pm-2.30am -0.052139250 -0.11793039  0.013651887 0.1582820
8.30am-2.30am -0.076666438 -0.14245757 -0.010875301 0.0176715*
8.30pm-2.30am -0.061616125 -0.12740726  0.004175012 0.0726070
8.30am-2.30pm -0.024527188 -0.09031832  0.041263949 0.7404688
8.30pm-2.30pm -0.009476875 -0.07526801  0.056314262 0.9789540
8.30pm-8.30am  0.015050313 -0.05074082  0.080841449 0.9233096

#10^3 concentration

            Df  Sum Sq  Mean Sq F value Pr(>F)
Time         3 0.01874 0.006248    1.41  0.261
Residuals   28 0.12405 0.004430

#10^2 concentration

            Df Sum Sq Mean Sq F value Pr(>F)
Time         3 0.1032 0.03440   3.766 0.0218 *
Residuals   28 0.2558 0.00914
---
Signif. codes:  0 ‘***’ 0.001 ‘**’ 0.01 ‘*’ 0.05 ‘.’ 0.1 ‘ ’ 1


Tukey multiple comparisons of means
    95% family-wise confidence level

$Time
                     diff         lwr         upr     p adj
2.30pm-2.30am -0.15597000 -0.28645430 -0.02548570 0.0145001*
8.30am-2.30am -0.10849581 -0.23898012  0.02198849 0.1294852
8.30pm-2.30am -0.07565313 -0.20613743  0.05483118 0.4042005
8.30am-2.30pm  0.04747419 -0.08301012  0.17795849 0.7543552
8.30pm-2.30pm  0.08031687 -0.05016743  0.21080118 0.3524424
8.30pm-8.30am  0.03284269 -0.09764162  0.16332699 0.9011016

#10^1 concentration

            Df Sum Sq Mean Sq F value Pr(>F)
Time         3 0.0925 0.03082   1.749   0.18
Residuals   28 0.4934 0.01762

#E2Nonenal

#10^5 concentration

            Df  Sum Sq  Mean Sq F value Pr(>F)
Time         3 0.02339 0.007796    1.67  0.196
Residuals   28 0.13074 0.004669

#10^4 concentration

            Df  Sum Sq  Mean Sq F value Pr(>F)
Time         3 0.01900 0.006333   2.783 0.0593 .
Residuals   28 0.06373 0.002276

#10^3 concentration

            Df  Sum Sq  Mean Sq F value Pr(>F)
Time         3 0.08062 0.026874   4.171 0.0146 *
Residuals   28 0.18043 0.006444
---
Signif. codes:  0 ‘***’ 0.001 ‘**’ 0.01 ‘*’ 0.05 ‘.’ 0.1 ‘ ’ 1

Tukey multiple comparisons of means
    95% family-wise confidence level

$Time
                     diff         lwr          upr     p adj
2.30pm-2.30am -0.12814976 -0.23773592 -0.018563609 0.0172032*
8.30am-2.30am -0.11041400 -0.22000015 -0.000827846 0.0477772*
8.30pm-2.30am -0.05543238 -0.16501853  0.054153779 0.5212043
8.30am-2.30pm  0.01773576 -0.09185039  0.127321916 0.9706374
8.30pm-2.30pm  0.07271739 -0.03686877  0.182303541 0.2891646
8.30pm-8.30am  0.05498163 -0.05460453  0.164567779 0.5279921


#10^2 concentration

            Df Sum Sq Mean Sq F value Pr(>F)
Time         3 0.0431 0.01437   1.077  0.375
Residuals   28 0.3737 0.01335

#10^1 concentration

            Df  Sum Sq  Mean Sq F value Pr(>F)
Time         3 0.04126 0.013753    1.45   0.25
Residuals   28 0.26563 0.009487

#Nonanal

#10^5 concentration

            Df  Sum Sq  Mean Sq F value Pr(>F)
Time         3 0.01961 0.006537   2.008  0.136
Residuals   28 0.09115 0.003255

#10^4 concentration

            Df  Sum Sq  Mean Sq F value Pr(>F)
Time         3 0.00917 0.003056   0.765  0.523
Residuals   28 0.11186 0.003995

#10^3 concentration

            Df  Sum Sq  Mean Sq F value Pr(>F)
Time         3 0.03684 0.012279   3.235 0.0371 *
Residuals   28 0.10628 0.003796
---
Signif. codes:  0 ‘***’ 0.001 ‘**’ 0.01 ‘*’ 0.05 ‘.’ 0.1 ‘ ’ 1

  Tukey multiple comparisons of means
    95% family-wise confidence level

$Time
                     diff         lwr          upr     p adj
2.30pm-2.30am -0.06505844 -0.14916442  0.019047540 0.1740101
8.30am-2.30am -0.09294600 -0.17705198 -0.008840022 0.0260740*
8.30pm-2.30am -0.04412254 -0.12822851  0.039983442 0.4905150
8.30am-2.30pm -0.02788756 -0.11199354  0.056218415 0.8021266
8.30pm-2.30pm  0.02093590 -0.06317008  0.105041880 0.9039478
8.30pm-8.30am  0.04882346 -0.03528251  0.132929442 0.4031338

#10^2 concentration

            Df Sum Sq Mean Sq F value   Pr(>F)
Time         3 0.1023 0.03411   7.499 0.000781 ***
Residuals   28 0.1274 0.00455
---
Signif. codes:  0 ‘***’ 0.001 ‘**’ 0.01 ‘*’ 0.05 ‘.’ 0.1 ‘ ’ 1

  Tukey multiple comparisons of means
    95% family-wise confidence level

$Time
                      diff           lwr         upr     p adj
2.30pm-2.30am -0.134289188 -0.2263659572 -0.04221242 0.0023540 **
8.30am-2.30am -0.126392000 -0.2184687697 -0.03431523 0.0043173 **
8.30pm-2.30am -0.043138786 -0.1352155554  0.04893798 0.5833456
8.30am-2.30pm  0.007897187 -0.0841795822  0.09997396 0.9953717
8.30pm-2.30pm  0.091150402 -0.0009263679  0.18322717 0.0531036
8.30pm-8.30am  0.083253214 -0.0088235554  0.17532998 0.0872506

#10^1 concentration

            Df Sum Sq Mean Sq F value   Pr(>F)
Time         3 0.1349 0.04496   9.852 0.000133 ***
Residuals   28 0.1278 0.00456
---
Signif. codes:  0 ‘***’ 0.001 ‘**’ 0.01 ‘*’ 0.05 ‘.’ 0.1 ‘ ’ 1

  Tukey multiple comparisons of means
    95% family-wise confidence level

$Time
                     diff         lwr         upr     p adj
2.30pm-2.30am -0.16841306 -0.26063262 -0.07619350 0.0001612 ***
8.30am-2.30am -0.12954400 -0.22176356 -0.03732444 0.0034461 **
8.30pm-2.30am -0.05756329 -0.14978285  0.03465628 0.3404668
8.30am-2.30pm  0.03886906 -0.05335050  0.13108862 0.6619158
8.30pm-2.30pm  0.11084978  0.01863022  0.20306934 0.0138690 *
8.30pm-8.30am  0.07198071 -0.02023885  0.16420028 0.1680643

#Benzaldehyde

#10^5 concentration

            Df  Sum Sq  Mean Sq F value  Pr(>F)
Time         3 0.02853 0.009508   4.648 0.00928 **
Residuals   28 0.05728 0.002046
---
Signif. codes:  0 ‘***’ 0.001 ‘**’ 0.01 ‘*’ 0.05 ‘.’ 0.1 ‘ ’ 1

  Tukey multiple comparisons of means
    95% family-wise confidence level

$Time
                      diff         lwr          upr     p adj
2.30pm-2.30am  0.008106156 -0.05364182  0.069854134 0.9838974
8.30am-2.30am -0.067546594 -0.12929457 -0.005798616 0.0279968*
8.30pm-2.30am -0.032201438 -0.09394942  0.029546540 0.4955602
8.30am-2.30pm -0.075652750 -0.13740073 -0.013904772 0.0118817*
8.30pm-2.30pm -0.040307594 -0.10205557  0.021440384 0.3027092
8.30pm-8.30am  0.035345156 -0.02640282  0.097093134 0.4153286


#10^4 concentration

            Df  Sum Sq  Mean Sq F value Pr(>F)
Time         3 0.04053 0.013510   3.791 0.0212 *
Residuals   28 0.09978 0.003564
---
Signif. codes:  0 ‘***’ 0.001 ‘**’ 0.01 ‘*’ 0.05 ‘.’ 0.1 ‘ ’ 1

  Tukey multiple comparisons of means
    95% family-wise confidence level

$Time
                      diff         lwr         upr     p adj
2.30pm-2.30am -0.036329094 -0.11782367  0.04516549 0.6214338
8.30am-2.30am -0.099055469 -0.18055005 -0.01756089 0.0126787*
8.30pm-2.30am -0.037446813 -0.11894139  0.04404777 0.5984486
8.30am-2.30pm -0.062726375 -0.14422096  0.01876821 0.1773273
8.30pm-2.30pm -0.001117719 -0.08261230  0.08037686 0.9999807
8.30pm-8.30am  0.061608656 -0.01988592  0.14310324 0.1896071

#10^3 concentration

            Df  Sum Sq  Mean Sq F value Pr(>F)
Time         3 0.02721 0.009069   2.071  0.127
Residuals   28 0.12261 0.004379

#10^2 concentration

            Df  Sum Sq  Mean Sq F value Pr(>F)
Time         3 0.04340 0.014466   4.163 0.0147 *
Residuals   28 0.09729 0.003475
---
Signif. codes:  0 ‘***’ 0.001 ‘**’ 0.01 ‘*’ 0.05 ‘.’ 0.1 ‘ ’ 1

  Tukey multiple comparisons of means
    95% family-wise confidence level

$Time
                     diff         lwr         upr     p adj
2.30pm-2.30am -0.10085109 -0.18132157 -0.02038062 0.0098335**
8.30am-2.30am -0.06991684 -0.15038732  0.01055363 0.1060393
8.30pm-2.30am -0.04620844 -0.12667891  0.03426204 0.4125723
8.30am-2.30pm  0.03093425 -0.04953622  0.11140472 0.7221756
8.30pm-2.30pm  0.05464266 -0.02582782  0.13511313 0.2704290
8.30pm-8.30am  0.02370841 -0.05676207  0.10417888 0.8517845

#10^1 concentration

            Df  Sum Sq  Mean Sq F value Pr(>F)
Time         3 0.02844 0.009479   1.619  0.207
Residuals   28 0.16393 0.005855

#Sulcatone

#10^5 concentration

            Df  Sum Sq  Mean Sq F value Pr(>F)
Time         3 0.00845 0.002817   1.901  0.152
Residuals   28 0.04148 0.001481

#10^4 concentration

            Df  Sum Sq  Mean Sq F value Pr(>F)
Time         3 0.00017 0.000057   0.046  0.987
Residuals   28 0.03504 0.001251


#10^3 concentration

            Df  Sum Sq  Mean Sq F value Pr(>F)
Time         3 0.00834 0.002781   1.182  0.334
Residuals   28 0.06587 0.002352

#10^2 concentration

            Df  Sum Sq  Mean Sq F value Pr(>F)
Time         3 0.03693 0.012309   2.461 0.0833 .
Residuals   28 0.14003 0.005001
---
Signif. codes:  0 ‘***’ 0.001 ‘**’ 0.01 ‘*’ 0.05 ‘.’ 0.1 ‘ ’ 1

#10^1 concentration

            Df  Sum Sq  Mean Sq F value Pr(>F)
Time         3 0.04364 0.014547   2.321 0.0968 .
Residuals   28 0.17546 0.006267
---
Signif. codes:  0 ‘***’ 0.001 ‘**’ 0.01 ‘*’ 0.05 ‘.’ 0.1 ‘ ’ 1

#3Octanol

#10^5 concentration

            Df  Sum Sq   Mean Sq F value Pr(>F)
Time         3 0.00105 0.0003505   0.239  0.868
Residuals   28 0.04100 0.0014645

#10^4 concentration

            Df  Sum Sq  Mean Sq F value Pr(>F)
Time         3 0.01154 0.003847   2.135  0.118
Residuals   28 0.05046 0.001802

#10^3 concentration

            Df  Sum Sq  Mean Sq F value Pr(>F)
Time         3 0.02238 0.007460   3.858 0.0199 *
Residuals   28 0.05414 0.001934
---
Signif. codes:  0 ‘***’ 0.001 ‘**’ 0.01 ‘*’ 0.05 ‘.’ 0.1 ‘ ’ 1

  Tukey multiple comparisons of means
    95% family-wise confidence level

$Time
                      diff         lwr          upr     p adj

2.30pm-2.30am -0.059426019 -0.11945761 0.0006055772 0.0431121*
8.30am-2.30am -0.056151506 -0.11618310 0.0038800897 0.0730989
8.30pm-2.30am -0.011056419 -0.07108801 0.0489751772 0.9577226
8.30am-2.30pm  0.003274512 -0.05675708 0.0633061084 0.9987933
8.30pm-2.30pm  0.048369600 -0.01166200 0.1084011959 0.1479823
8.30pm-8.30am  0.045095088 -0.01493651 0.1051266834 0.1940493

#10^2 concentration

            Df  Sum Sq  Mean Sq F value Pr(>F)
Time         3 0.04997 0.016658   3.793 0.0212 *
Residuals   28 0.12297 0.004392
---
Signif. codes:  0 ‘***’ 0.001 ‘**’ 0.01 ‘*’ 0.05 ‘.’ 0.1 ‘ ’ 1

  Tukey multiple comparisons of means
    95% family-wise confidence level


$Time
                      diff         lwr         upr     p adj
2.30pm-2.30am -0.108003794 -0.19847306 -0.01753453 0.0146448 *
8.30am-2.30am -0.037577256 -0.12804652  0.05289201 0.6720268
8.30pm-2.30am -0.030845294 -0.12131456  0.05962397 0.7886263
8.30am-2.30pm  0.070426537 -0.02004273  0.16089580 0.1698110
8.30pm-2.30pm  0.077158500 -0.01331076  0.16762776 0.1155951
8.30pm-8.30am  0.006731963 -0.08373730  0.09720123 0.9969605

#10^1 concentration

            Df  Sum Sq  Mean Sq F value Pr(>F)
Time         3 0.02674 0.008914    2.25  0.104
Residuals   28 0.11093 0.003962


Including species identity as a predictor
```

Results of repeated measure ANOVA for EAG read outs of all compounds using species, dose, and time as predictors where individual identity (ID) is nested in dose.

Model used: Volatile compound/blend ~ Species * Dose * Time + Error (ID/Dose)

```
Camphor

Error: ID
             Df Sum Sq  Mean Sq F value Pr(>F)
Species       1 0.0017 0.001690   0.111  0.740
Time          3 0.0303 0.010085   0.663  0.578
Species:Time  3 0.0251 0.008354   0.549  0.651
Residuals    56 0.8516 0.015206

Error: ID:Dose
                   Df Sum Sq Mean Sq F value   Pr(>F)
Dose                4 0.7045 0.17611  95.308  < 2e-16 ***
Species:Dose        4 0.1149 0.02873  15.546 3.07e-11 ***
Dose:Time          12 0.0221 0.00184   0.998   0.4520
Species:Dose:Time  12 0.0385 0.00321   1.735   0.0607 .
Residuals         224 0.4139 0.00185
---
Signif. codes:  0 ‘***’ 0.001 ‘**’ 0.01 ‘*’ 0.05 ‘.’ 0.1 ‘ ’ 1


Geraniol

Error: ID
             Df Sum Sq Mean Sq F value   Pr(>F)
Species       1 0.7261  0.7261  27.283 2.68e-06 ***
Time          3 0.0510  0.0170   0.638    0.594
Species:Time  3 0.0283  0.0094   0.355    0.786
Residuals    56 1.4903  0.0266
---
Signif. codes:  0 ‘***’ 0.001 ‘**’ 0.01 ‘*’ 0.05 ‘.’ 0.1 ‘ ’ 1

Error: ID:Dose
                   Df Sum Sq Mean Sq F value   Pr(>F)
Dose                4  3.707  0.9268 210.585  < 2e-16 ***
Species:Dose        4  0.267  0.0667  15.147 5.64e-11 ***
Dose:Time          12  0.040  0.0033   0.760   0.6910
Species:Dose:Time  12  0.092  0.0077   1.744   0.0591 .
Residuals         224  0.986  0.0044
---
Signif. codes:  0 ‘***’ 0.001 ‘**’ 0.01 ‘*’ 0.05 ‘.’ 0.1 ‘ ’ 1

4MePhenol

Error: ID
             Df Sum Sq Mean Sq F value   Pr(>F)
Species       1 0.1184 0.11842  17.014 0.000124 ***
Time          3 0.0345 0.01149   1.652 0.187895
Species:Time  3 0.0435 0.01449   2.083 0.112830
Residuals    56 0.3898 0.00696
---
Signif. codes:  0 ‘***’ 0.001 ‘**’ 0.01 ‘*’ 0.05 ‘.’ 0.1 ‘ ’ 1

Error: ID:Dose
                   Df Sum Sq Mean Sq F value  Pr(>F)
Dose                4 0.3557 0.08892  88.201 < 2e-16 ***
Species:Dose        4 0.0780 0.01950  19.347 1.1e-13 ***
Dose:Time          12 0.0218 0.00181   1.799  0.0495 *
Species:Dose:Time  12 0.0140 0.00117   1.160  0.3135
Residuals         224 0.2258 0.00101
---
Signif. codes:  0 ‘***’ 0.001 ‘**’ 0.01 ‘*’ 0.05 ‘.’ 0.1 ‘ ’ 1

Acetophenone

Error: ID
             Df Sum Sq Mean Sq F value   Pr(>F)
Species       1  4.919   4.919 102.924 2.71e-14 ***
Time          3  0.147   0.049   1.023   0.3892
Species:Time  3  0.392   0.131   2.732   0.0522 .
Residuals    56  2.676   0.048
---
Signif. codes:  0 ‘***’ 0.001 ‘**’ 0.01 ‘*’ 0.05 ‘.’ 0.1 ‘ ’ 1

Error: ID:Dose
                   Df Sum Sq Mean Sq F value Pr(>F)
Dose                4  6.695  1.6736 165.638 <2e-16 ***
Species:Dose        4  1.028  0.2569  25.423 <2e-16 ***
Dose:Time          12  0.024  0.0020   0.194  0.999
Species:Dose:Time  12  0.150  0.0125   1.238  0.258
Residuals         224  2.263  0.0101
---
Signif. codes:  0 ‘***’ 0.001 ‘**’ 0.01 ‘*’ 0.05 ‘.’ 0.1 ‘ ’ 1

Phenol

Error: ID
             Df Sum Sq Mean Sq F value   Pr(>F)
Species       1 0.1267 0.12672  19.894 3.99e-05 ***
Time          3 0.0086 0.00288   0.452   0.7166
Species:Time  3 0.0494 0.01647   2.586   0.0621 .
Residuals    56 0.3567 0.00637
---
Signif. codes:  0 ‘***’ 0.001 ‘**’ 0.01 ‘*’ 0.05 ‘.’ 0.1 ‘ ’ 1

Error: ID:Dose
                   Df  Sum Sq  Mean Sq F value  Pr(>F)
Dose                4 0.06550 0.016374  25.862 < 2e-16 ***
Species:Dose        4 0.01129 0.002821   4.456 0.00175 **
Dose:Time          12 0.02152 0.001794   2.833 0.00122 **
Species:Dose:Time  12 0.01195 0.000996   1.572 0.10091
Residuals         224 0.14182 0.000633
---
Signif. codes:  0 ‘***’ 0.001 ‘**’ 0.01 ‘*’ 0.05 ‘.’ 0.1 ‘ ’ 1

β Ocimine

Error: ID
             Df Sum Sq Mean Sq F value   Pr(>F)
Species       1 0.8380  0.8380  27.795 2.24e-06 ***
Time          3 0.0949  0.0316   1.049    0.378
Species:Time  3 0.1587  0.0529   1.755    0.166
Residuals    56 1.6883  0.0301
---
Signif. codes:  0 ‘***’ 0.001 ‘**’ 0.01 ‘*’ 0.05 ‘.’ 0.1 ‘ ’ 1

Error: ID:Dose
                   Df Sum Sq Mean Sq F value   Pr(>F)
Dose                4  9.009  2.2524 183.493  < 2e-16 ***
Species:Dose        4  0.703  0.1757  14.313 2.02e-10 ***
Dose:Time          12  0.231  0.0192   1.568  0.10235
Species:Dose:Time  12  0.335  0.0279   2.272  0.00976 **
Residuals         224  2.750  0.0123
---
Signif. codes:  0 ‘***’ 0.001 ‘**’ 0.01 ‘*’ 0.05 ‘.’ 0.1 ‘ ’ 1

E2Nonenal

Error: ID
             Df Sum Sq Mean Sq F value Pr(>F)
Species       1 0.1405 0.14047   5.671 0.0207 *
Time          3 0.1422 0.04740   1.914 0.1379
Species:Time  3 0.0829 0.02763   1.116 0.3505
Residuals    56 1.3871 0.02477
---
Signif. codes:  0 ‘***’ 0.001 ‘**’ 0.01 ‘*’ 0.05 ‘.’ 0.1 ‘ ’ 1

Error: ID:Dose
                   Df Sum Sq Mean Sq F value Pr(>F)
Dose                4 2.1252  0.5313 112.654 <2e-16 ***
Species:Dose        4 0.0401  0.0100   2.128 0.0783 .
Dose:Time          12 0.0309  0.0026   0.546 0.8830
Species:Dose:Time  12 0.0461  0.0038   0.815 0.6352
Residuals         224 1.0565  0.0047
---
Signif. codes:  0 ‘***’ 0.001 ‘**’ 0.01 ‘*’ 0.05 ‘.’ 0.1 ‘ ’ 1

Benzaldehyde


Error: ID
             Df Sum Sq Mean Sq F value   Pr(>F)
Species       1  8.297   8.297 121.356 1.22e-15 ***
Time          3  0.242   0.081   1.182    0.325
Species:Time  3  0.097   0.032   0.475    0.701
Residuals    56  3.829   0.068
---
Signif. codes:  0 ‘***’ 0.001 ‘**’ 0.01 ‘*’ 0.05 ‘.’ 0.1 ‘ ’ 1

Error: ID:Dose
                   Df Sum Sq Mean Sq F value Pr(>F)
Dose                4 11.840  2.9600 176.529 <2e-16 ***
Species:Dose        4  3.250  0.8125  48.458 <2e-16 ***
Dose:Time          12  0.199  0.0166   0.989  0.460
Species:Dose:Time  12  0.199  0.0166   0.991  0.458
Residuals         224  3.756  0.0168
---
Signif. codes:  0 ‘***’ 0.001 ‘**’ 0.01 ‘*’ 0.05 ‘.’ 0.1 ‘ ’ 1

Nonanal

Error: ID
             Df Sum Sq Mean Sq F value   Pr(>F)
Species       1 1.1639  1.1639  26.646 3.34e-06 ***
Time          3 0.1005  0.0335   0.767    0.517
Species:Time  3 0.1962  0.0654   1.498    0.225
Residuals    56 2.4460  0.0437
---
Signif. codes:  0 ‘***’ 0.001 ‘**’ 0.01 ‘*’ 0.05 ‘.’ 0.1 ‘ ’ 1

Error: ID:Dose
                   Df Sum Sq Mean Sq F value Pr(>F)
Dose                4  9.120  2.2799 210.468 <2e-16 ***
Species:Dose        4  1.271  0.3177  29.326 <2e-16 ***
Dose:Time          12  0.062  0.0052   0.479 0.9256
Species:Dose:Time  12  0.278  0.0232   2.140 0.0156 *
Residuals         224  2.426  0.0108
---
Signif. codes:  0 ‘***’ 0.001 ‘**’ 0.01 ‘*’ 0.05 ‘.’ 0.1 ‘ ’ 1

Limonene

Error: ID
             Df Sum Sq Mean Sq F value   Pr(>F)
Species       1  3.268   3.268  61.669 1.35e-10 ***
Time          3  0.092   0.031   0.576    0.633
Species:Time  3  0.064   0.021   0.402    0.752
Residuals    56  2.968   0.053
---
Signif. codes:  0 ‘***’ 0.001 ‘**’ 0.01 ‘*’ 0.05 ‘.’ 0.1 ‘ ’ 1

Error: ID:Dose
                   Df Sum Sq Mean Sq F value Pr(>F)
Dose                4  2.766  0.6916  44.228 <2e-16 ***
Species:Dose        4  1.513  0.3782  24.188 <2e-16 ***
Dose:Time          12  0.155  0.0129   0.827  0.623
Species:Dose:Time  12  0.126  0.0105   0.674  0.776
Residuals         224  3.503  0.0156
---
Signif. codes:  0 ‘***’ 0.001 ‘**’ 0.01 ‘*’ 0.05 ‘.’ 0.1 ‘ ’ 1

Sulcatone

Error: ID
             Df Sum Sq Mean Sq F value   Pr(>F)
Species       1  3.543   3.543   82.90 1.22e-12 ***
Time          3  0.204   0.068    1.59    0.202
Species:Time  3  0.173   0.058    1.35    0.268
Residuals    56  2.393   0.043
---
Signif. codes:  0 ‘***’ 0.001 ‘**’ 0.01 ‘*’ 0.05 ‘.’ 0.1 ‘ ’ 1

Error: ID:Dose
                   Df Sum Sq Mean Sq F value   Pr(>F)
Dose                4  8.066  2.0165 125.968  < 2e-16 ***
Species:Dose        4  1.424  0.3560  22.237 1.78e-15 ***
Dose:Time          12  0.079  0.0066   0.412    0.958
Species:Dose:Time  12  0.112  0.0093   0.581    0.856
Residuals         224  3.586  0.0160
---
Signif. codes:  0 ‘***’ 0.001 ‘**’ 0.01 ‘*’ 0.05 ‘.’ 0.1 ‘ ’ 1

3Octanol

Error: ID
             Df Sum Sq Mean Sq F value   Pr(>F)
Species       1 2.8997  2.8997 116.599 2.63e-15 ***
Time          3 0.1639  0.0546   2.196   0.0986 .
Species:Time  3 0.2888  0.0963   3.871   0.0138 *
Residuals    56 1.3927  0.0249
---
Signif. codes:  0 ‘***’ 0.001 ‘**’ 0.01 ‘*’ 0.05 ‘.’ 0.1 ‘ ’ 1

Error: ID:Dose
                   Df Sum Sq Mean Sq F value Pr(>F)
Dose                4  6.401  1.6003 255.223 <2e-16 ***
Species:Dose        4  1.417  0.3543  56.507 <2e-16 ***
Dose:Time          12  0.091  0.0076   1.210  0.277
Species:Dose:Time  12  0.088  0.0073   1.165  0.310
Residuals         224  1.405  0.0063
---
Signif. codes:  0 ‘***’ 0.001 ‘**’ 0.01 ‘*’ 0.05 ‘.’ 0.1 ‘ ’ 1


Human Odor Blend

Error: ID
             Df Sum Sq Mean Sq F value Pr(>F)
Species       1 15.199  15.199 150.961 <2e-16 ***
Time          3  0.269   0.090   0.891  0.451
Species:Time  3  0.295   0.098   0.978  0.410
Residuals    56  5.638   0.101
---
Signif. codes:  0 ‘***’ 0.001 ‘**’ 0.01 ‘*’ 0.05 ‘.’ 0.1 ‘ ’ 1

Error: ID:Dose
                   Df Sum Sq Mean Sq F value Pr(>F)
Dose                4 15.593   3.898 209.077 <2e-16 ***
Species:Dose        4  6.551   1.638  87.841 <2e-16 ***
Dose:Time          12  0.198   0.017   0.887  0.561
Species:Dose:Time  12  0.294   0.025   1.316  0.210
Residuals         224  4.176   0.019
---
Signif. codes:  0 ‘***’ 0.001 ‘**’ 0.01 ‘*’ 0.05 ‘.’ 0.1 ‘ ’ 1
```

Results of two-way ANOVA and post hoc Tukey HSD test for the EAG readouts for 4MePhenol, Phenol, Ocimene, Nonanal and 3Octanol to figure out the difference between different time points at each concentration/dose. Compounds showing differences between the time points were highlighted by blue, concentration by green and time points pair by grey.

```
#4MePhenol

#10^5 concentration

             Df  Sum Sq  Mean Sq F value  Pr(>F)
Species       1 0.00005 0.000055   0.044 0.83510
Time          3 0.00889 0.002963   2.366 0.08065 .
Species:Time  3 0.01908 0.006361   5.078 0.00352 **
Residuals    56 0.07014 0.001252
---
Signif. codes:  0 ‘***’ 0.001 ‘**’ 0.01 ‘*’ 0.05 ‘.’ 0.1 ‘ ’ 1

  Tukey multiple comparisons of means
    95% family-wise confidence level


$`Species:Time`
                                           diff          lwr          upr     p adj
Anopheles:2.30am-Aedes:2.30am     -0.0414366875 -0.097146423  0.014273048 0.2902836
Aedes:2.30pm-Aedes:2.30am         -0.0118463750 -0.067556111  0.043863361 0.9974818
Anopheles:2.30pm-Aedes:2.30am     -0.0344856250 -0.090195361  0.021224111 0.5245057
Aedes:8.30am-Aedes:2.30am         -0.0656654500 -0.121375186 -0.009955714 0.0105197
Anopheles:8.30am-Aedes:2.30am     -0.0301106250 -0.085820361  0.025599111 0.6861526
Aedes:8.30pm-Aedes:2.30am         -0.0367908125 -0.092500548  0.018918923 0.4405931
Anopheles:8.30pm-Aedes:2.30am     -0.0008680625 -0.056577798  0.054841673 1.0000000
Aedes:2.30pm-Anopheles:2.30am      0.0295903125 -0.026119423  0.085300048 0.7045689
Anopheles:2.30pm-Anopheles:2.30am  0.0069510625 -0.048758673  0.062660798 0.9999238
Aedes:8.30am-Anopheles:2.30am     -0.0242287625 -0.079938498  0.031480973 0.8671014
Anopheles:8.30am-Anopheles:2.30am  0.0113260625 -0.044383673  0.067035798 0.9981047
Aedes:8.30pm-Anopheles:2.30am      0.0046458750 -0.051063861  0.060355611 0.9999951
Anopheles:8.30pm-Anopheles:2.30am  0.0405686250 -0.015141111  0.096278361 0.3158577
Anopheles:2.30pm-Aedes:2.30pm     -0.0226392500 -0.078348986  0.033070486 0.9026326
Aedes:8.30am-Aedes:2.30pm         -0.0538190750 -0.109528811  0.001890661 0.0654186
Anopheles:8.30am-Aedes:2.30pm     -0.0182642500 -0.073973986  0.037445486 0.9673803
Aedes:8.30pm-Aedes:2.30pm         -0.0249444375 -0.080654173  0.030765298 0.8490057
Anopheles:8.30pm-Aedes:2.30pm      0.0109783125 -0.044731423  0.066688048 0.9984464
Aedes:8.30am-Anopheles:2.30pm     -0.0311798250 -0.086889561  0.024529911 0.6474460
Anopheles:8.30am-Anopheles:2.30pm  0.0043750000 -0.051334736  0.060084736 0.9999968
Aedes:8.30pm-Anopheles:2.30pm     -0.0023051875 -0.058014923  0.053404548 1.0000000
Anopheles:8.30pm-Anopheles:2.30pm  0.0336175625 -0.022092173  0.089327298 0.5568436
Anopheles:8.30am-Aedes:8.30am      0.0355548250 -0.020154911  0.091264561 0.4851072
Aedes:8.30pm-Aedes:8.30am          0.0288746375 -0.026835098  0.084584373 0.7293361
Anopheles:8.30pm-Aedes:8.30am      0.0647973875  0.009087652  0.120507123 0.0121553
Aedes:8.30pm-Anopheles:8.30am     -0.0066801875 -0.062389923  0.049029548 0.9999418
Anopheles:8.30pm-Anopheles:8.30am  0.0292425625 -0.026467173  0.084952298 0.7166911
Anopheles:8.30pm-Aedes:8.30pm      0.0359227500 -0.019786986  0.091632486 0.4717200


#10^4 concentration

            Df  Sum Sq  Mean Sq F value  Pr(>F)
Species       1 0.00335 0.003351   2.499 0.11955
Time          3 0.00250 0.000832   0.620 0.60469
Species:Time  3 0.01769 0.005898   4.399 0.00755 **
Residuals    56 0.07509 0.001341
---
Signif. codes:  0 ‘***’ 0.001 ‘**’ 0.01 ‘*’ 0.05 ‘.’ 0.1 ‘ ’ 1

  Tukey multiple comparisons of means
    95% family-wise confidence level


$`Species:Time`
                                          diff          lwr        upr     p adj
Anopheles:2.30am-Aedes:2.30am     -0.039291063 -0.096931980 0.01834985 0.3994180
Aedes:2.30pm-Aedes:2.30am         -0.018700913 -0.076341830 0.03894000 0.9691845
Anopheles:2.30pm-Aedes:2.30am     -0.004513625 -0.062154542 0.05312729 0.9999968
Aedes:8.30am-Aedes:2.30am         -0.044371500 -0.102012417 0.01326942 0.2505459
Anopheles:8.30am-Aedes:2.30am     -0.008888625 -0.066529542 0.04875229 0.9996857
Aedes:8.30pm-Aedes:2.30am         -0.035479438 -0.093120355 0.02216148 0.5317465
Anopheles:8.30pm-Aedes:2.30am      0.012027937 -0.045612980 0.06966885 0.9977644
Aedes:2.30pm-Anopheles:2.30am      0.020590150 -0.037050767 0.07823107 0.9485860
Anopheles:2.30pm-Anopheles:2.30am  0.034777438 -0.022863480 0.09241835 0.5570414
Aedes:8.30am-Anopheles:2.30am     -0.005080437 -0.062721355 0.05256048 0.9999929
Anopheles:8.30am-Anopheles:2.30am  0.030402438 -0.027238480 0.08804335 0.7117853
Aedes:8.30pm-Anopheles:2.30am      0.003811625 -0.053829292 0.06145254 0.9999990
Anopheles:8.30pm-Anopheles:2.30am  0.051319000 -0.006321917 0.10895992 0.1149086
Anopheles:2.30pm-Aedes:2.30pm      0.014187288 -0.043453630 0.07182820 0.9937734
Aedes:8.30am-Aedes:2.30pm         -0.025670587 -0.083311505 0.03197033 0.8524879
Anopheles:8.30am-Aedes:2.30pm      0.009812288 -0.047828630 0.06745320 0.9993974
Aedes:8.30pm-Aedes:2.30pm         -0.016778525 -0.074419442 0.04086239 0.9832144
Anopheles:8.30pm-Aedes:2.30pm      0.030728850 -0.026912067 0.08836977 0.7007371
Aedes:8.30am-Anopheles:2.30pm     -0.039857875 -0.097498792 0.01778304 0.3809523
Anopheles:8.30am-Anopheles:2.30pm -0.004375000 -0.062015917 0.05326592 0.9999975
Aedes:8.30pm-Anopheles:2.30pm     -0.030965813 -0.088606730 0.02667510 0.6926401
Anopheles:8.30pm-Anopheles:2.30pm  0.016541562 -0.041099355 0.07418248 0.9845327
Anopheles:8.30am-Aedes:8.30am      0.035482875 -0.022158042 0.09312379 0.5316230
Aedes:8.30pm-Aedes:8.30am          0.008892062 -0.048748855 0.06653298 0.9996849
Anopheles:8.30pm-Aedes:8.30am      0.056399437 -0.001241480 0.11404035 0.0593661
Aedes:8.30pm-Anopheles:8.30am     -0.026590813 -0.084231730 0.03105010 0.8285521
Anopheles:8.30pm-Anopheles:8.30am  0.020916562 -0.036724355 0.07855748 0.9442511
Anopheles:8.30pm-Aedes:8.30pm      0.047507375 -0.010133542 0.10514829 0.1796180

#10^3 concentration

             Df  Sum Sq  Mean Sq F value   Pr(>F)
Species       1 0.02352 0.023518  12.109 0.000979 ***
Time          3 0.00609 0.002029   1.045 0.379927
Species:Time  3 0.00884 0.002948   1.518 0.219889
Residuals    56 0.10876 0.001942
---
Signif. codes:  0 ‘***’ 0.001 ‘**’ 0.01 ‘*’ 0.05 ‘.’ 0.1 ‘ ’ 1

#10^2 concentration

             Df  Sum Sq Mean Sq F value  Pr(>F)
Species       1 0.03279 0.03279  15.035 0.00028 ***
Time          3 0.00305 0.00102   0.466 0.70725
Species:Time  3 0.00983 0.00328   1.502 0.22395
Residuals    56 0.12213 0.00218
---
Signif. codes:  0 ‘***’ 0.001 ‘**’ 0.01 ‘*’ 0.05 ‘.’ 0.1 ‘ ’ 1

#10^1 concentration

            Df  Sum Sq Mean Sq F value   Pr(>F)
Species       1 0.13673 0.13673  31.973 5.51e-07 ***
Time          3 0.03572 0.01191   2.785   0.0491 *
Species:Time  3 0.00207 0.00069   0.161   0.9219
Residuals    56 0.23948 0.00428
---
Signif. codes:  0 ‘***’ 0.001 ‘**’ 0.01 ‘*’ 0.05 ‘.’ 0.1 ‘ ’ 1

  Tukey multiple comparisons of means
    95% family-wise confidence level

$`Species:Time`
                                          diff          lwr          upr     p adj
Anopheles:2.30am-Aedes:2.30am      0.094551312 -0.008387818  0.197490443 0.0937454
Aedes:2.30pm-Aedes:2.30am         -0.034227625 -0.137166756  0.068711506 0.9647940
Anopheles:2.30pm-Aedes:2.30am      0.039670250 -0.063268881  0.142609381 0.9245997
Aedes:8.30am-Aedes:2.30am         -0.064964875 -0.167904006  0.037974256 0.4996383
Anopheles:8.30am-Aedes:2.30am      0.031545250 -0.071393881  0.134484381 0.9775295
Aedes:8.30pm-Aedes:2.30am         -0.030949438 -0.133888568  0.071989693 0.9798181
Anopheles:8.30pm-Aedes:2.30am      0.073856562 -0.029082568  0.176795693 0.3342111
Aedes:2.30pm-Anopheles:2.30am     -0.128778937 -0.231718068 -0.025839807 0.0052802
Anopheles:2.30pm-Anopheles:2.30am -0.054881062 -0.157820193  0.048058068 0.7006731
Aedes:8.30am-Anopheles:2.30am     -0.159516187 -0.262455318 -0.056577057 0.0002363
Anopheles:8.30am-Anopheles:2.30am -0.063006062 -0.165945193  0.039933068 0.5389091
Aedes:8.30pm-Anopheles:2.30am     -0.125500750 -0.228439881 -0.022561619 0.0071791
Anopheles:8.30pm-Anopheles:2.30am -0.020694750 -0.123633881  0.082244381 0.9982352
Anopheles:2.30pm-Aedes:2.30pm      0.073897875 -0.029041256  0.176837006 0.3335146
Aedes:8.30am-Aedes:2.30pm         -0.030737250 -0.133676381  0.072201881 0.9805896
Anopheles:8.30am-Aedes:2.30pm      0.065772875 -0.037166256  0.168712006 0.4836152
Aedes:8.30pm-Aedes:2.30pm          0.003278187 -0.099660943  0.106217318 1.0000000
Anopheles:8.30pm-Aedes:2.30pm      0.108084187  0.005145057  0.211023318 0.0330901
Aedes:8.30am-Anopheles:2.30pm     -0.104635125 -0.207574256 -0.001695994 0.0437338
Anopheles:8.30am-Anopheles:2.30pm -0.008125000 -0.111064131  0.094814131 0.9999967
Aedes:8.30pm-Anopheles:2.30pm     -0.070619688 -0.173558818  0.032319443 0.3911415
Anopheles:8.30pm-Anopheles:2.30pm  0.034186312 -0.068752818  0.137125443 0.9650225
Anopheles:8.30am-Aedes:8.30am      0.096510125 -0.006429006  0.199449256 0.0813710
Aedes:8.30pm-Aedes:8.30am          0.034015437 -0.068923693  0.136954568 0.9659563
Anopheles:8.30pm-Aedes:8.30am      0.138821438  0.035882307  0.241760568 0.0019945
Aedes:8.30pm-Anopheles:8.30am     -0.062494688 -0.165433818  0.040444443 0.5492277
Anopheles:8.30pm-Anopheles:8.30am  0.042311312 -0.060627818  0.145250443 0.8972837
Anopheles:8.30pm-Aedes:8.30pm      0.104806000  0.001866869  0.207745131 0.0431426

#Phenol

#10^5 concentration

             Df  Sum Sq Mean Sq F value  Pr(>F)
Species       1 0.04264 0.04264  17.106 0.00012 ***
Time          3 0.00814 0.00271   1.088 0.36163
Species:Time  3 0.01700 0.00567   2.273 0.09001 .
Residuals    56 0.13958 0.00249
---
Signif. codes:  0 ‘***’ 0.001 ‘**’ 0.01 ‘*’ 0.05 ‘.’ 0.1 ‘ ’ 1

#10^4 concentration

             Df  Sum Sq  Mean Sq F value   Pr(>F)
Species       1 0.02789 0.027887  17.044 0.000123 ***
Time          3 0.00583 0.001943   1.187 0.322883
Species:Time  3 0.01571 0.005238   3.202 0.030068 *
Residuals    56 0.09162 0.001636
---
Signif. codes:  0 ‘***’ 0.001 ‘**’ 0.01 ‘*’ 0.05 ‘.’ 0.1 ‘ ’ 1

  Tukey multiple comparisons of means
    95% family-wise confidence level


$`Species:Time`
                                           diff         lwr          upr     p adj
Anopheles:2.30am-Aedes:2.30am     -0.0879953125 -0.15166769 -0.024322934 0.0014143
Aedes:2.30pm-Aedes:2.30am         -0.0105352500 -0.07420763  0.053137129 0.9994998
Anopheles:2.30pm-Aedes:2.30am     -0.0425085000 -0.10618088  0.021163879 0.4264407
Aedes:8.30am-Aedes:2.30am         -0.0504317232 -0.11410410  0.013240656 0.2195184
Anopheles:8.30am-Aedes:2.30am     -0.0512585000 -0.11493088  0.012413879 0.2027063
Aedes:8.30pm-Aedes:2.30am         -0.0092238125 -0.07289619  0.054448566 0.9997925
Anopheles:8.30pm-Aedes:2.30am     -0.0554218125 -0.11909419  0.008250566 0.1320536
Aedes:2.30pm-Anopheles:2.30am      0.0774600625  0.01378768  0.141132441 0.0073619
Anopheles:2.30pm-Anopheles:2.30am  0.0454868125 -0.01818557  0.109159191 0.3395996
Aedes:8.30am-Anopheles:2.30am      0.0375635893 -0.02610879  0.101235968 0.5849235
Anopheles:8.30am-Anopheles:2.30am  0.0367368125 -0.02693557  0.100409191 0.6118857
Aedes:8.30pm-Anopheles:2.30am      0.0787715000  0.01509912  0.142443879 0.0060410
Anopheles:8.30pm-Anopheles:2.30am  0.0325735000 -0.03109888  0.096245879 0.7420059
Anopheles:2.30pm-Aedes:2.30pm     -0.0319732500 -0.09564563  0.031699129 0.7593831
Aedes:8.30am-Aedes:2.30pm         -0.0398964732 -0.10356885  0.023775906 0.5088979
Anopheles:8.30am-Aedes:2.30pm     -0.0407232500 -0.10439563  0.022949129 0.4823438
Aedes:8.30pm-Aedes:2.30pm          0.0013114375 -0.06236094  0.064983816 1.0000000
Anopheles:8.30pm-Aedes:2.30pm     -0.0448865625 -0.10855894  0.018785816 0.3563270
Aedes:8.30am-Anopheles:2.30pm     -0.0079232232 -0.07159560  0.055749156 0.9999252
Anopheles:8.30am-Anopheles:2.30pm -0.0087500000 -0.07242238  0.054922379 0.9998541
Aedes:8.30pm-Anopheles:2.30pm      0.0332846875 -0.03038769  0.096957066 0.7208482
Anopheles:8.30pm-Anopheles:2.30pm -0.0129133125 -0.07658569  0.050759066 0.9981339
Anopheles:8.30am-Aedes:8.30am     -0.0008267768 -0.06449916  0.062845602 1.0000000
Aedes:8.30pm-Aedes:8.30am          0.0412079107 -0.02246447  0.104880290 0.4669488
Anopheles:8.30pm-Aedes:8.30am     -0.0049900893 -0.06866247  0.058682290 0.9999968
Aedes:8.30pm-Anopheles:8.30am      0.0420346875 -0.02163769  0.105707066 0.4410481
Anopheles:8.30pm-Anopheles:8.30am -0.0041633125 -0.06783569  0.059509066 0.9999991
Anopheles:8.30pm-Aedes:8.30pm     -0.0461980000 -0.10987038  0.017474379 0.3203441


#10^3 concentration

             Df  Sum Sq Mean Sq F value   Pr(>F)
Species       1 0.04337 0.04337  24.505 7.16e-06 ***
Time          3 0.00627 0.00209   1.181   0.3251
Species:Time  3 0.01368 0.00456   2.576   0.0629 .
Residuals    56 0.09912 0.00177
---
Signif. codes:  0 ‘***’ 0.001 ‘**’ 0.01 ‘*’ 0.05 ‘.’ 0.1 ‘ ’ 1


#10^2 concentration

             Df  Sum Sq  Mean Sq F value  Pr(>F)
Species       1 0.01721 0.017215  11.726 0.00116 **
Time          3 0.00266 0.000888   0.605 0.61469
Species:Time  3 0.01005 0.003351   2.282 0.08900 .
Residuals    56 0.08221 0.001468
---
Signif. codes:  0 ‘***’ 0.001 ‘**’ 0.01 ‘*’ 0.05 ‘.’ 0.1 ‘ ’ 1


#10^1 concentration

             Df  Sum Sq  Mean Sq F value Pr(>F)
Species       1 0.00690 0.006898   4.492 0.0385 *
Time          3 0.00727 0.002422   1.577 0.2051
Species:Time  3 0.00492 0.001641   1.069 0.3698
Residuals    56 0.08600 0.001536
---
Signif. codes:  0 ‘***’ 0.001 ‘**’ 0.01 ‘*’ 0.05 ‘.’ 0.1 ‘ ’ 1

#Ocimene

#10^5 concentration

             Df  Sum Sq  Mean Sq F value Pr(>F)
Species       1 0.00109 0.001088   0.239  0.627
Time          3 0.01094 0.003646   0.801  0.499
Species:Time  3 0.00085 0.000283   0.062  0.980
Residuals    56 0.25507 0.004555


#10^4 concentration

             Df Sum Sq  Mean Sq F value Pr(>F)
Species       1 0.0141 0.014084   2.147 0.1484
Time          3 0.0497 0.016561   2.525 0.0668 .
Species:Time  3 0.0022 0.000730   0.111 0.9531
Residuals    56 0.3673 0.006559
---
Signif. codes:  0 ‘***’ 0.001 ‘**’ 0.01 ‘*’ 0.05 ‘.’ 0.1 ‘ ’ 1

#10^3 concentration

            Df Sum Sq Mean Sq F value Pr(>F)
Species       1 0.1061 0.10611   5.558 0.0219 *
Time          3 0.0535 0.01785   0.935 0.4300
Species:Time  3 0.0071 0.00237   0.124 0.9453
Residuals    56 1.0692 0.01909
---
Signif. codes:  0 ‘***’ 0.001 ‘**’ 0.01 ‘*’ 0.05 ‘.’ 0.1 ‘ ’ 1

#10^2 concentration

             Df Sum Sq Mean Sq F value  Pr(>F)
Species       1 0.2288 0.22878  10.073 0.00244 **
Time          3 0.1068 0.03561   1.568 0.20730
Species:Time  3 0.0801 0.02671   1.176 0.32717
Residuals    56 1.2719 0.02271
---
Signif. codes:  0 ‘***’ 0.001 ‘**’ 0.01 ‘*’ 0.05 ‘.’ 0.1 ‘ ’ 1

#10^1 concentration

             Df Sum Sq Mean Sq F value   Pr(>F)
Species       1 1.1907  1.1907  45.221 9.86e-09 ***
Time          3 0.1048  0.0349   1.327  0.27488
Species:Time  3 0.4031  0.1344   5.103  0.00342 **
Residuals    56 1.4745  0.0263
---
Signif. codes:  0 ‘***’ 0.001 ‘**’ 0.01 ‘*’ 0.05 ‘.’ 0.1 ‘ ’ 1

Tukey multiple comparisons of means
    95% family-wise confidence level
$`Species:Time`
                                          diff         lwr          upr     p adj
Anopheles:2.30am-Aedes:2.30am      0.063433937 -0.19199336  0.318861233 0.9934262
Aedes:2.30pm-Aedes:2.30am         -0.128581875 -0.38400917  0.126845421 0.7571123
Anopheles:2.30pm-Aedes:2.30am      0.373776000  0.11834870  0.629203296 0.0006011
Aedes:8.30am-Aedes:2.30am         -0.132576063 -0.38800336  0.122851233 0.7279455
Anopheles:8.30am-Aedes:2.30am      0.175026000 -0.08040130  0.430453296 0.3926567
Aedes:8.30pm-Aedes:2.30am         -0.071480750 -0.32690805  0.183946546 0.9866305
Anopheles:8.30pm-Aedes:2.30am      0.146304000 -0.10912330  0.401731296 0.6205398
Aedes:2.30pm-Anopheles:2.30am     -0.192015812 -0.44744311  0.063411483 0.2777404
Anopheles:2.30pm-Anopheles:2.30am  0.310342063  0.05491477  0.565769358 0.0074715
Aedes:8.30am-Anopheles:2.30am     -0.196010000 -0.45143730  0.059417296 0.2540737
Anopheles:8.30am-Anopheles:2.30am  0.111592063 -0.14383523  0.367019358 0.8644050
Aedes:8.30pm-Anopheles:2.30am     -0.134914687 -0.39034198  0.120512608 0.7103358
Anopheles:8.30pm-Anopheles:2.30am  0.082870063 -0.17255723  0.338297358 0.9691851
Anopheles:2.30pm-Aedes:2.30pm      0.502357875  0.24693058  0.757785171 0.0000020
Aedes:8.30am-Aedes:2.30pm         -0.003994187 -0.25942148  0.251433108 1.0000000
Anopheles:8.30am-Aedes:2.30pm      0.303607875  0.04818058  0.559035171 0.0095877
Aedes:8.30pm-Aedes:2.30pm          0.057101125 -0.19832617  0.312528421 0.9965554
Anopheles:8.30pm-Aedes:2.30pm      0.274885875  0.01945858  0.530313171 0.0264502
Aedes:8.30am-Anopheles:2.30pm     -0.506352062 -0.76177936 -0.250924767 0.0000017
Anopheles:8.30am-Anopheles:2.30pm -0.198750000 -0.45417730  0.056677296 0.2386271
Aedes:8.30pm-Anopheles:2.30pm     -0.445256750 -0.70068405 -0.189829454 0.0000269
Anopheles:8.30pm-Anopheles:2.30pm -0.227472000 -0.48289930  0.027955296 0.1147195
Anopheles:8.30am-Aedes:8.30am      0.307602062  0.05217477  0.563029358 0.0082735
Aedes:8.30pm-Aedes:8.30am          0.061095312 -0.19433198  0.316522608 0.9947704
Anopheles:8.30pm-Aedes:8.30am      0.278880062  0.02345277  0.534307358 0.0230858
Aedes:8.30pm-Anopheles:8.30am     -0.246506750 -0.50193405  0.008920546 0.0659220
Anopheles:8.30pm-Anopheles:8.30am -0.028722000 -0.28414930  0.226705296 0.9999623
Anopheles:8.30pm-Aedes:8.30pm      0.217784750 -0.03764255  0.473212046 0.1490394

#Nonanal

#10^5 concentration

             Df Sum Sq  Mean Sq F value Pr(>F)
Species       1 0.0044 0.004428   0.439  0.510
Time          3 0.0407 0.013553   1.344  0.269
Species:Time  3 0.0051 0.001689   0.168  0.918
Residuals    56 0.5646 0.010082


#10^4 concentration

            Df Sum Sq  Mean Sq F value Pr(>F)
Species       1 0.0058 0.005823   0.529  0.470
Time          3 0.0123 0.004107   0.373  0.773
Species:Time  3 0.0011 0.000368   0.033  0.992
Residuals    56 0.6167 0.011013

#10^3 concentration

             Df Sum Sq Mean Sq F value Pr(>F)
Species       1 0.0624 0.06242   2.450  0.123
Time          3 0.0238 0.00794   0.312  0.817
Species:Time  3 0.0400 0.01333   0.523  0.668
Residuals    56 1.4266 0.02547


#10^2 concentration

             Df Sum Sq Mean Sq F value   Pr(>F)
Species       1 1.1036  1.1036  42.196 2.35e-08 ***
Time          3 0.0078  0.0026   0.099    0.960
Species:Time  3 0.2134  0.0711   2.720    0.053 .
Residuals    56 1.4646  0.0262
---
Signif. codes:  0 ‘***’ 0.001 ‘**’ 0.01 ‘*’ 0.05 ‘.’ 0.1 ‘ ’ 1

#10^1 concentration

Df Sum Sq Mean Sq F value   Pr(>F)
Species       1 1.2583  1.2583  88.087 4.32e-13 ***
Time          3 0.0782  0.0261   1.825  0.15312
Species:Time  3 0.2148  0.0716   5.012  0.00378 **
Residuals    56 0.8000  0.0143
---
Signif. codes:  0 ‘***’ 0.001 ‘**’ 0.01 ‘*’ 0.05 ‘.’ 0.1 ‘ ’ 1

Tukey multiple comparisons of means
    95% family-wise confidence level

$`Species:Time`
                                         diff         lwr          upr     p adj
Anopheles:2.30am-Aedes:2.30am      0.21237187  0.02423007  0.400513684 0.0166241
Aedes:2.30pm-Aedes:2.30am         -0.16841306 -0.35655487  0.019728747 0.1110408
Anopheles:2.30pm-Aedes:2.30am      0.27507425  0.08693244  0.463216059 0.0006093
Aedes:8.30am-Aedes:2.30am         -0.12954400 -0.31768581  0.058597809 0.3864334
Anopheles:8.30am-Aedes:2.30am      0.19819925  0.01005744  0.386341059 0.0321308
Aedes:8.30pm-Aedes:2.30am         -0.05756329 -0.24570510  0.130578524 0.9777299
Anopheles:8.30pm-Aedes:2.30am      0.08058637 -0.10755543  0.268728184 0.8758759
Aedes:2.30pm-Anopheles:2.30am     -0.38078494 -0.56892675 -0.192643128 0.0000010
Anopheles:2.30pm-Anopheles:2.30am  0.06270238 -0.12543943  0.250844184 0.9643536
Aedes:8.30am-Anopheles:2.30am     -0.34191587 -0.53005768 -0.153774066 0.0000115
Anopheles:8.30am-Anopheles:2.30am -0.01417262 -0.20231443  0.173969184 0.9999976
Aedes:8.30pm-Anopheles:2.30am     -0.26993516 -0.45807697 -0.081793351 0.0008144
Anopheles:8.30pm-Anopheles:2.30am -0.13178550 -0.31992731  0.056356309 0.3644647
Anopheles:2.30pm-Aedes:2.30pm      0.44348731  0.25534550  0.631629122 0.0000000
Aedes:8.30am-Aedes:2.30pm          0.03886906 -0.14927275  0.227010872 0.9979015
Anopheles:8.30am-Aedes:2.30pm      0.36661231  0.17847050  0.554754122 0.0000025
Aedes:8.30pm-Aedes:2.30pm          0.11084978 -0.07729203  0.298991586 0.5865235
Anopheles:8.30pm-Aedes:2.30pm      0.24899944  0.06085763  0.437141247 0.0025738
Aedes:8.30am-Anopheles:2.30pm     -0.40461825 -0.59276006 -0.216476441 0.0000002
Anopheles:8.30am-Anopheles:2.30pm -0.07687500 -0.26501681  0.111266809 0.9000997
Aedes:8.30pm-Anopheles:2.30pm     -0.33263754 -0.52077935 -0.144495726 0.0000203
Anopheles:8.30pm-Anopheles:2.30pm -0.19448788 -0.38262968 -0.006346066 0.0379240
Anopheles:8.30am-Aedes:8.30am      0.32774325  0.13960144  0.515885059 0.0000273
Aedes:8.30pm-Aedes:8.30am          0.07198071 -0.11616110  0.260122524 0.9272502
Anopheles:8.30pm-Aedes:8.30am      0.21013037  0.02198857  0.398272184 0.0184984
Aedes:8.30pm-Anopheles:8.30am     -0.25576254 -0.44390435 -0.067620726 0.0017851
Anopheles:8.30pm-Anopheles:8.30am -0.11761288 -0.30575468  0.070528934 0.5119051
Anopheles:8.30pm-Aedes:8.30pm      0.13814966 -0.04999215  0.326291470 0.3057421

#3Octanol

#10^5 concentration

             Df Sum Sq Mean Sq F value Pr(>F)
Species       1 0.0521 0.05206   6.888 0.0112 *
Time          3 0.0025 0.00082   0.109 0.9546
Species:Time  3 0.0046 0.00152   0.202 0.8947
Residuals    56 0.4233 0.00756
---
Signif. codes:  0 ‘***’ 0.001 ‘**’ 0.01 ‘*’ 0.05 ‘.’ 0.1 ‘ ’ 1

#10^4 concentration

            Df Sum Sq  Mean Sq F value Pr(>F)
Species       1 0.0268 0.026779   2.672  0.108
Time          3 0.0245 0.008167   0.815  0.491
Species:Time  3 0.0268 0.008944   0.893  0.451
Residuals    56 0.5612 0.010021

#10^3 concentration

            Df Sum Sq Mean Sq F value  Pr(>F)
Species       1 0.4310  0.4310  46.289 7.3e-09 ***
Time          3 0.0213  0.0071   0.762  0.5201
Species:Time  3 0.0623  0.0208   2.229  0.0948 .
Residuals    56 0.5214  0.0093
---
Signif. codes:  0 ‘***’ 0.001 ‘**’ 0.01 ‘*’ 0.05 ‘.’ 0.1 ‘ ’ 1

#10^2 concentration

            Df Sum Sq Mean Sq F value  Pr(>F)
Species       1 1.9137  1.9137 156.853 < 2e-16 ***
Time          3 0.1145  0.0382   3.128 0.03277 *
Species:Time  3 0.1598  0.0533   4.366 0.00784 **
Residuals    56 0.6832  0.0122
---
Signif. codes:  0 ‘***’ 0.001 ‘**’ 0.01 ‘*’ 0.05 ‘.’ 0.1 ‘ ’ 1

  Tukey multiple comparisons of means
    95% family-wise confidence level

$`Species:Time`
                                          diff         lwr           upr     p adj
Anopheles:2.30am-Aedes:2.30am      0.317533563  0.14366027  0.4914068568 0.0000103
Aedes:2.30pm-Aedes:2.30am         -0.108003794 -0.28187709  0.0658695005 0.5200823
Anopheles:2.30pm-Aedes:2.30am      0.337166500  0.16329321  0.5110397943 0.0000028
Aedes:8.30am-Aedes:2.30am         -0.037577256 -0.21145055  0.1362960380 0.9972117
Anopheles:8.30am-Aedes:2.30am      0.388416500  0.21454321  0.5622897943 0.0000001
Aedes:8.30pm-Aedes:2.30am         -0.030845294 -0.20471859  0.1430280005 0.9992113
Anopheles:8.30pm-Aedes:2.30am      0.163819250 -0.01005404  0.3376925443 0.0785767
Aedes:2.30pm-Anopheles:2.30am     -0.425537356 -0.59941065 -0.2516640620 0.0000000
Anopheles:2.30pm-Anopheles:2.30am  0.019632938 -0.15424036  0.1935062318 0.9999613
Aedes:8.30am-Anopheles:2.30am     -0.355110819 -0.52898411 -0.1812375245 0.0000008
Anopheles:8.30am-Anopheles:2.30am  0.070882937 -0.10299036  0.2447562318 0.9011659
Aedes:8.30pm-Anopheles:2.30am     -0.348378856 -0.52225215 -0.1745055620 0.0000013
Anopheles:8.30pm-Anopheles:2.30am -0.153714313 -0.32758761  0.0201589818 0.1200997
Anopheles:2.30pm-Aedes:2.30pm      0.445170294  0.27129700  0.6190435880 0.0000000
Aedes:8.30am-Aedes:2.30pm          0.070426537 -0.10344676  0.2442998318 0.9041331
Anopheles:8.30am-Aedes:2.30pm      0.496420294  0.32254700  0.6702935880 0.0000000
Aedes:8.30pm-Aedes:2.30pm          0.077158500 -0.09671479  0.2510317943 0.8547682
Anopheles:8.30pm-Aedes:2.30pm      0.271823044  0.09794975  0.4456963380 0.0002032
Aedes:8.30am-Anopheles:2.30pm     -0.374743756 -0.54861705 -0.2008704620 0.0000002
Anopheles:8.30am-Anopheles:2.30pm  0.051250000 -0.12262329  0.2251232943 0.9819681
Aedes:8.30pm-Anopheles:2.30pm     -0.368011794 -0.54188509 -0.1941384995 0.0000003
Anopheles:8.30pm-Anopheles:2.30pm -0.173347250 -0.34722054  0.0005260443 0.0512324
Anopheles:8.30am-Aedes:8.30am      0.425993756  0.25212046  0.5998670505 0.0000000
Aedes:8.30pm-Aedes:8.30am          0.006731963 -0.16714133  0.1806052568 1.0000000
Anopheles:8.30pm-Aedes:8.30am      0.201396506  0.02752321  0.3752698005 0.0127083
Aedes:8.30pm-Anopheles:8.30am     -0.419261794 -0.59313509 -0.2453884995 0.0000000
Anopheles:8.30pm-Anopheles:8.30am -0.224597250 -0.39847054 -0.0507239557 0.0035374
Anopheles:8.30pm-Aedes:8.30pm      0.194664544  0.02079125  0.3685378380 0.0180567

#10^1 concentration

    Df Sum Sq Mean Sq F value Pr(>F)
Species       1 1.8935  1.8935 174.350 <2e-16 ***
Time          3 0.0922  0.0307   2.828 0.0466 *
Species:Time  3 0.1230  0.0410   3.774 0.0154 *
Residuals    56 0.6082  0.0109
---
Signif. codes:  0 ‘***’ 0.001 ‘**’ 0.01 ‘*’ 0.05 ‘.’ 0.1 ‘ ’ 1

  Tukey multiple comparisons of means
    95% family-wise confidence level


$`Species:Time`
                                          diff         lwr          upr     p adj
Anopheles:2.30am-Aedes:2.30am      0.327695312  0.16364839  0.491742238 0.0000014
Aedes:2.30pm-Aedes:2.30am         -0.076055544 -0.24010247  0.087991381 0.8249466
Anopheles:2.30pm-Aedes:2.30am      0.347139875  0.18309295  0.511186800 0.0000003
Aedes:8.30am-Aedes:2.30am         -0.018369631 -0.18241656  0.145677294 0.9999634
Anopheles:8.30am-Aedes:2.30am      0.399639875  0.23559295  0.563686800 0.0000000
Aedes:8.30pm-Aedes:2.30am         -0.015437669 -0.17948459  0.148609256 0.9999888
Anopheles:8.30pm-Aedes:2.30am      0.191714875  0.02766795  0.355761800 0.0115492
Aedes:2.30pm-Anopheles:2.30am     -0.403750856 -0.56779778 -0.239703931 0.0000000
Anopheles:2.30pm-Anopheles:2.30am  0.019444563 -0.14460236  0.183491488 0.9999462
Aedes:8.30am-Anopheles:2.30am     -0.346064944 -0.51011187 -0.182018019 0.0000004
Anopheles:8.30am-Anopheles:2.30am  0.071944563 -0.09210236  0.235991488 0.8620894
Aedes:8.30pm-Anopheles:2.30am     -0.343132981 -0.50717991 -0.179086056 0.0000005
Anopheles:8.30pm-Anopheles:2.30am -0.135980437 -0.30002736  0.028066488 0.1742450
Anopheles:2.30pm-Aedes:2.30pm      0.423195419  0.25914849  0.587242344 0.0000000
Aedes:8.30am-Aedes:2.30pm          0.057685912 -0.10636101  0.221732838 0.9526193
Anopheles:8.30am-Aedes:2.30pm      0.475695419  0.31164849  0.639742344 0.0000000
Aedes:8.30pm-Aedes:2.30pm          0.060617875 -0.10342905  0.224664800 0.9388608
Anopheles:8.30pm-Aedes:2.30pm      0.267770419  0.10372349  0.431817344 0.0000945
Aedes:8.30am-Anopheles:2.30pm     -0.365509506 -0.52955643 -0.201462581 0.0000001
Anopheles:8.30am-Anopheles:2.30pm  0.052500000 -0.11154693  0.216546925 0.9714091
Aedes:8.30pm-Anopheles:2.30pm     -0.362577544 -0.52662447 -0.198530619 0.0000001
Anopheles:8.30pm-Anopheles:2.30pm -0.155425000 -0.31947193  0.008621925 0.0754976
Anopheles:8.30am-Aedes:8.30am      0.418009506  0.25396258  0.582056431 0.0000000
Aedes:8.30pm-Aedes:8.30am          0.002931962 -0.16111496  0.166978888 1.0000000
Anopheles:8.30pm-Aedes:8.30am      0.210084506  0.04603758  0.374131431 0.0039487
Aedes:8.30pm-Anopheles:8.30am     -0.415077544 -0.57912447 -0.251030619 0.0000000
Anopheles:8.30pm-Anopheles:8.30am -0.207925000 -0.37197193 -0.043878075 0.0044956
Anopheles:8.30pm-Aedes:8.30pm      0.207152544  0.04310562  0.371199469 0.0047081
```
